# Dynein-microtubule forces drive nucleokinesis and transmigration in T cells

**DOI:** 10.64898/2026.07.02.736211

**Authors:** Yerbol Tagay, Alexander S. Zhovmer, Nandini Sarkar, Jorik Stoop, Linjun Su, Laurel Fleszar, Eric Peterman, Jeffrey P. Rasmussen, Alexander X. Cartagena-Rivera, Denis Tsygankov, Erdem D. Tabdanov

**Affiliations:** Department of Cell and Biological Systems, Penn State College of Medicine, The Pennsylvania State University, Hershey, PA, USA; Section on Mechanobiology, National Institute of Biomedical Imaging and Bioengineering, National Institutes of Health, Bethesda, MD, USA; Center for Biologics Evaluation and Research, U.S. Food and Drug Administration, Silver Spring, MD, USA; Wallace H. Coulter Department of Biomedical Engineering, Georgia Institute of Technology and Emory University, Atlanta, GA, USA; Department of Biology, University of Washington, Seattle, WA, USA; Institute for Stem Cell and Regenerative Medicine, University of Washington, Seattle, WA, USA; Penn State Cancer Institute, Penn State College of Medicine, The Pennsylvania State University, Hershey, PA, USA

**Keywords:** T Lymphocyte, Uropod, Septin, Amoeboid Motility, Biophysical Simulations, Cell Confinement

## Abstract

Beyond the mechanical capacity of canonical actomyosin-driven amoeboid motility in permissive extracellular environments, the nucleus becomes the principal barrier to T cell migration in confining tissues. We establish the dynein-microtubule (MT) force-transmission axis as an essential mechanism for nuclear translocation during confined T cell migration and transmigration. We argue that dynein acts both as a motor and as an F-actin-anchored force-transmission element (fulcrum), sliding MTs and the MT-coupled nucleus along the cell cortex to drive nucleokinesis and productive cell displacement. Dynein is the primary driver of nucleokinesis: its inhibition arrests nucleus movement independently of myosin II activity, while F-actin dynamics remain spatiotemporally decoupled from nuclear oscillations. During transmigration, dynein and actomyosin act cooperatively and non-redundantly, and only combined inhibition abolishes nuclear passage. Computational modeling demonstrates that dynein-mediated pulling, together with volume exclusion imposed by the nucleus, is sufficient to generate self-organized nuclear oscillations. Dynein inhibition in zebrafish Langerhans cells impairs protrusion dynamics *in situ*, identifying the dynein-MT axis as an evolutionarily conserved mechanobiological program. Collectively, these findings identify the dynein-MT mechanical unit as a potential target for engineering T cells with enhanced solid-tumor infiltration.

## INTRODUCTION

Cell migration in densely packed solid tissues, as well as the associated cellular morphological rearrangements, are characteristic features of immune surveillance ^1–4^, neural plasticity ^5–7^, and metastatic motility ^8–11^. For cells navigating through narrow interstitial spaces, the near-continuous confinement imposed by densely packed three-dimensional tissues fundamentally differs from the crowding imposed by loosely structured, discrete fibrillar obstacles, such as collagen, in extracellular matrices (ECM) and stromal regions^12–14^. Whereas fibrillar obstacles predominantly induce amoeboid migration in cells and cytoplasts ^15–18^, continuous or overly dense confinement promotes mesenchymal behavior that relies on cell adhesion ^19,20^, coordinated force generation ^21–23^, and long-distance force transmission ^14,23–26^ along the confined ^27,28^, elongated, and/or branched (dendritic-like) arborization of the cell body ^29–31^. Understanding how cells sense and adapt to environmental constraints is therefore central to understanding their capacity for deep-tissue infiltration, including into solid tumors.

Recent reports have highlighted the intact microtubule (MT) network as an essential structure that spatiotemporally regulates and integrates actomyosin contractility signaling, thereby enabling mechanically coherent migration under confinement. This coordination allows net cell displacement without excessive cell-body extension or rupture ^27,28,32–34^. In migrating immune cells subjected to confinement-induced narrowing, elongation, and overextension, the microtubule-organizing center (MTOC) and its associated dense MT network help define the cell’s leading protrusion. Conversely, the MT-deficient rear engages the GEF-H1→RhoA→non-muscle myosin II pathway: the local absence of MTs prevents sequestration of GEF-H1 ^35^, thereby promoting actomyosin contractility and generation of trailing forces that define the cell’s rear ^27^. In partially confined cancer cells, rear MTs compressed between the advancing nucleus and the trailing contractile actomyosin cortex undergo mechanical breakage and depolymerization, releasing GEF-H1 and further amplifying RhoA-myosin II-dependent contractility at the collapsing rear. Thus, MTs may function as a mechanostat whose mechanical fragility sets the deformation threshold that drives nuclear translocation and cell movement ^36^. However, both mechanisms emphasize the signaling functions of MTs and do not address their potential role as direct mediators of force generation and transmission during cell migration.

Evidence increasingly supports a more direct mechanical role for MTs and MT-associated motors in migratory force generation and transmission ^37–43^. For instance, dynein can structurally and mechanically integrate into the actin cytoskeleton through dynactin, a dynein-activating cofactor and adaptor complex ^44,45^. Because dynactin contains an F-actin-like Arp1 filament that can be primed to nucleate F-actin polymerization^46^, it may function as a structural bridge linking microtubule-based dynein forces to the actin network ^45–47^, associated integrin adhesion complexes, and ECM. Alternatively, dynein can anchor to F-actin *via* the Afadin-LGN-NuMA ^48^, ADAP ^49^, Spectrin-Dynactin ^50^, and Annexin-A2 ^51^ molecular adapter complexes. Thus, dynein anchored to F-actin has been proposed to generate propulsive forces that support cell locomotion in the absence of actomyosin contractility ^40^ or on mechanically compliant substrates, where mechanosensing-dependent actomyosin contractility is attenuated or absent ^41^. Structurally, as relatively inextensible load-bearing cables, microtubules could relay mechanical forces, including dynein-generated forces ^52–56^, over long distances within cells ^40,57^ and cell protrusions, such as axons ^58–60^, with minimal mechanical dissipation.

The nucleus is the largest and stiffest organelle, constituting a major structural and mechanical obstacle to migration through narrow interstitial spaces ^27,36,61–63^. Cells have evolved various actomyosin-driven strategies to move the nucleus in confinement, including mechanical ^36,64,65^ or hydraulic ^17,64,66–68^ nuclear piston mechanisms, and actomyosin-dependent nuclear pulling ^62,69^ or pushing ^67,70–72^ mechanisms. However, an alternative strategy relies on MT-associated nucleokinesis. During neocortex development, neuronal precursors move their nuclei tens of micrometers along glial fibers ^72–74^, using dynein-generated pulling forces to slide microtubules and nuclei ^6^. This process requires the MT-associated protein LIS1 ^75^, which facilitates dynein complex assembly ^76–78^ and anchors MTs to the F-actin cortex ^79^. Similarly, during skeletal myoblast-to-myotube fusion, nuclei are repositioned along syncytium microtubules through the coordinated activity of dynein/dynactin motor complexes and cell-polarity regulators ^80^. Together, these observations demonstrate that extensive mechanical coupling between the nucleus and the MT cytoskeleton can support long-range nuclear translocation, suggesting that similar mechanisms may contribute to nuclear propulsion during confined immune- and cancer-cell migration.

In summary, inefficient trafficking of cytotoxic lymphocytes into solid tumors remains a major barrier to anticancer immunity ^81–85^ due to: (**1**) dense collagen at the tumor-stroma interface ^81,86^, (**2**) abnormal tumor vasculature and elevated interstitial fluid pressure ^87^, and (**3**) confining tissue architecture with subnuclear-size interstitial spaces ^88^. While actomyosin-dependent amoeboid T cell motility is increasingly well-defined, studies in other cell types show that dynein-driven nuclear translocation along microtubule cables contributes to migration through dense tissues. This raises the possibility that a comparable dynein-MT-F-actin force-transmission mechanism could enable T cells to deform, polarize, and propel their nuclei through the crowded tumor interstitium. However, contributions of dynein and MTs as a long-range intracellular force-transmission system ^40,41^ remain poorly characterized in T cells. We therefore hypothesized that propulsion of the large and mechanically resistant nucleus, a major burden during T cell migration through confining tissue environments, requires coordinated generation and transmission of forces across the cytoskeletal network, including dynein and MTs. Elucidating the mechanism that drives nucleokinesis and transmigration may reveal a previously underappreciated axis of cytoskeletal cooperation governing T cell motility within solid tumors and provide a missing mechanobiological element for engineering immune cells optimized for deep-tissue penetration and durable antitumor activity.

## RESULTS

### A conceptual framework for amoeboid and mesenchymal T cell mechanotypes

The physical properties of the environment, particularly geometry and degree of confinement, impose distinct mechanical demands on migrating T cells, thereby selecting for distinct force-generating, force-sensing, and force-transmission strategies, *i.e.*, mechanotypes.

At one extreme, discrete crowding by fibrillar obstacles, such as individual collagen fibers in the extracellular matrix, induces the amoeboid mechanotype **(Figure 1a-1)**. In this mode, septins act as mechanosensors and “proprioceptors” that detect cell surface indentations caused by steric obstacles and template the formation of circumferential F-actin cortical actomyosin rings at those sites ^15,89^ **(Figure 1b)**. The resulting cortical rings partition the actomyosin cortex into hydraulically linked contractile compartments, enabling peristaltic cytosolic flow and pressure-driven piston-like nuclear propulsion that collectively power amoeboid locomotion through loosely structured environments ^15,17^. Consequently, peristaltically driven cytoplasmic flow ^15^ and the piston-like nucleus translocation ^17^ cooperate to propel amoeboid migration through confining interstitial spaces **(Figure 1a-1)**. Consistent with this model, primary human CD4^+^ T cells **(Figure 1b)** form F-actin rings at sites of interaction with collagen fibers **(Figure 1b)**, supporting a role for microenvironmental geometry in coordinating the adaptive reorganization of the amoeboid actomyosin cortex. Notably, pharmacological inhibition of septin GTPase activity disrupts septin filament assembly, abolishes cortical ring formation, and suppresses the amoeboid mechanotype and motility ^15^. Furthermore, while amoeboid migration is integrin-independent and does not require strong cell-ECM adhesion or long-range transmission of traction forces, its mechanical efficiency appears to be limited by the degree of confinement. For instance, amoeboid (bleb-driven) melanoma cells migrate through dense isotropic 3D collagen at only ∼0.027 µm/min ^25^, whereas mesenchymally migrating breast cancer cells in comparable matrices move approximately six times faster, at ∼0.167 µm/min ^90^.

**Figure 1.**
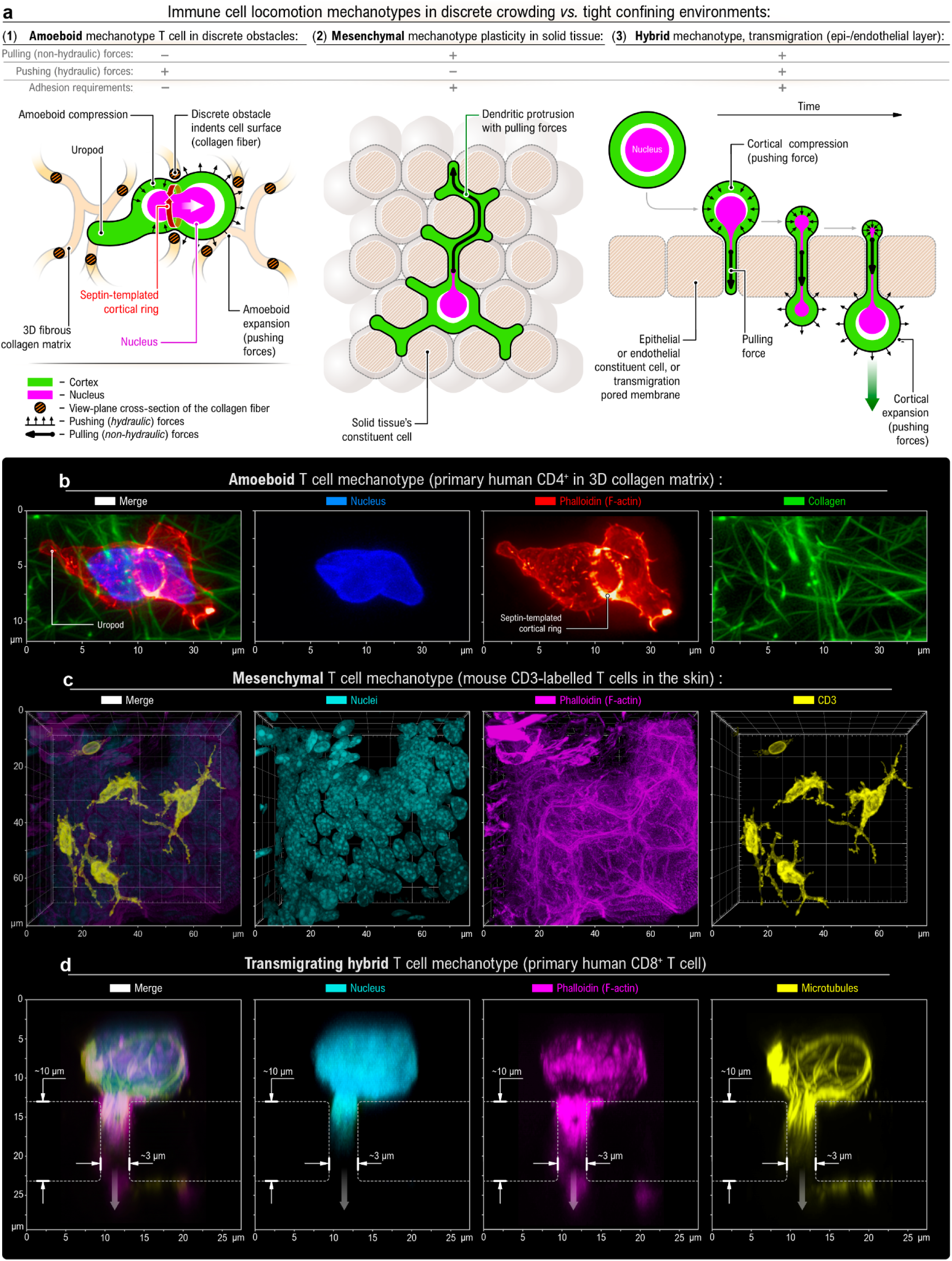
Environmental context determines T cell migration mechanotype across discrete, continuous, and barrier confinements. **a** - T cells deploy distinct migration mechanotypes (force generation and transmission mechanisms) in various microenvironments: **(1)** Discrete cell-crowding obstacles induce an amoeboid mechanotype: Individual collagen fibers crowd and indent the T cell surface. At the sites of indentation, septin triggers the formation of circumferential cortical rings. Multiple cytoskeletal rings partition the actomyosin cortex into contractile segments. Multiple hydraulically linked contractile compartments enable peristaltic translocation of the cytosol, *i.e.*, amoeboid motility. In this mode, cell propulsion is based on peristaltic, hydraulically driven pushing and expansion forces, independent of cell-ECM traction (pulling) forces and adhesion. **(2)** Continuous confinement within solid tissues induces a mesenchymal mechanotype. T cells extend thin dendritic protrusions between the tissue’s tightly packed constituent cells. Non-hydraulic traction (pulling) forces generated within these protrusions drive cell propulsion through adhesion-mediated force transmission. In contrast to amoeboid motility mechanisms, the mesenchymal-like locomotion mode requires strong cell-ECM adhesion and substantial traction-based pulling forces. **(3)** Hybrid propulsion during T cell migration across epithelial or endothelial barriers. During transmigration (diapedesis), T cells employ both pushing and pulling forces, either simultaneously or sequentially. Cortical compression generates hydraulic forces that help the cell wedge the nucleus between adjacent cells. Upon nuclear entry into the pore, the transmigrating cell uses traction or hydraulic forces to pull or push the nucleus through the intercellular gap. Upon exiting, the rear cortical compression pushed the nucleus out of the intercellular gap. **b** - Amoeboid mechanotype, featuring cortical-ring-mediated cell segmentation and an amoeboid polarity axis defined by the uropod, in a human primary CD4^+^ T cell within a 3D collagen type-I matrix. **c** - Mesenchymal mechanotype. 3D rendering of skin-resident mouse CD3⁺ T cells showing branched cell morphology conforming to the interstitial space between tissue constituent cells. **d** - Transmigration of human primary CD8^+^ T cells through a 3 µm-wide, 10 µm-long ICAM1-coated polyurethane pore.

At the other extreme, continuous confinement within densely packed solid tissues, where interstitial spaces are narrower than the cell body and often narrower than the nucleus itself ^88^, fundamentally differs from discrete fibrillar crowding ^12–14^ and promotes a mesenchymal-like mechanotype ^91,92^ **(Figure 1a-2)**. Here, “mesenchymal” refers to the mechanobiological state of T cells, defined by their morphology, adhesion, and force-transmission properties, rather than to their lineage identity.

**Figure 2.**
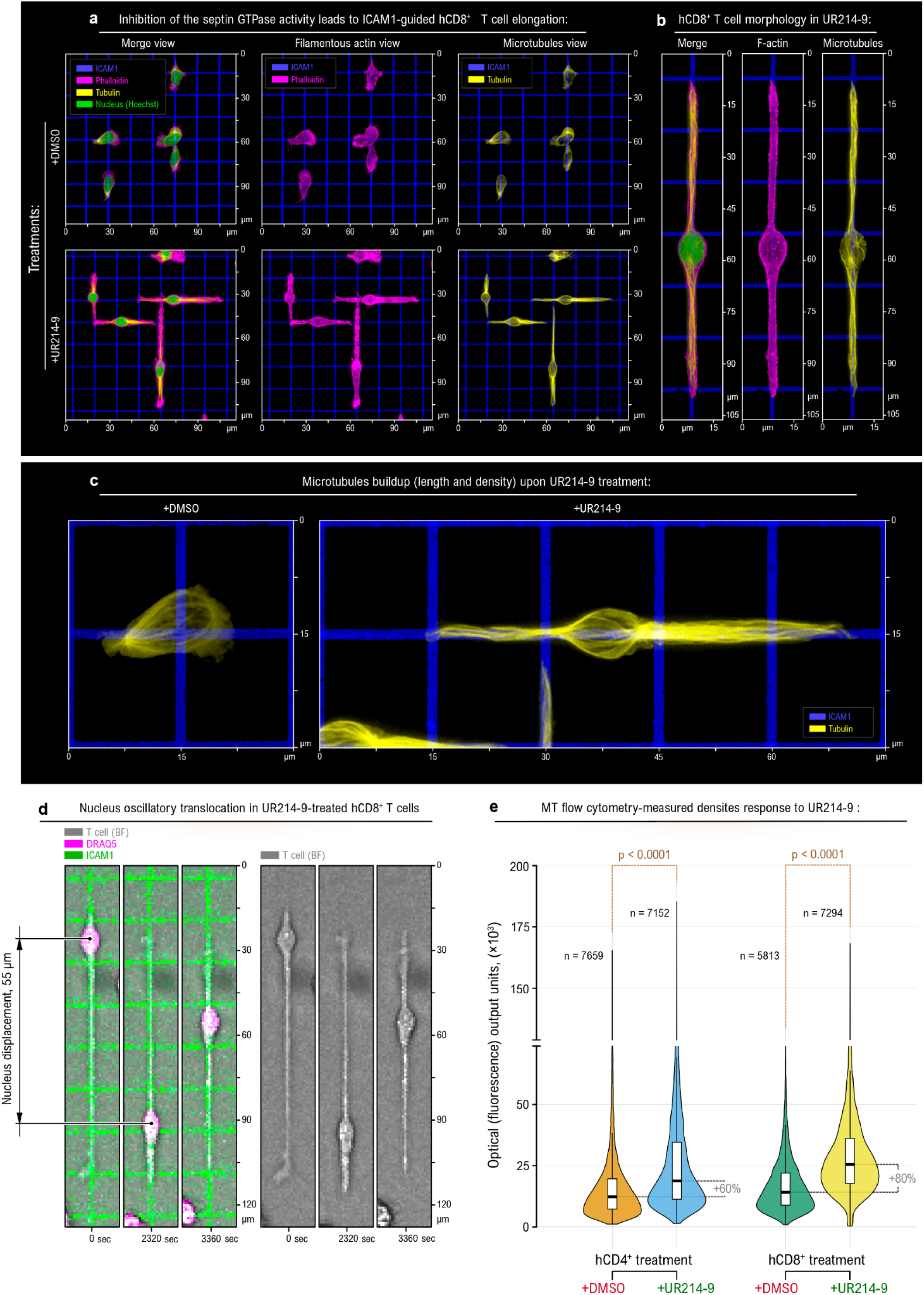
Septin inhibition drives a phenotypic amoeboid-to-mesenchymal transition, accompanied by expansion of the microtubule network, in primary hCD8^+^ T cells. **a** - *Top row* (+DMSO) - Non-treated control hCD8^+^ T cells display a mixed amoeboid-mesenchymal phenotype (*immunofluorescence, 3D reconstruction, top view*). Although lymphocytes adhere to the ICAM1 micropattern (blue), they maintain a predominantly amoeboid morphology, characterized by compact, non-spread shapes, poor alignment with the underlying adhesive micropattern’s geometry, and an underdeveloped microtubule network (*yellow*). *Bottom row* (+UR214-9) - Inhibition of septin GTPase activity with UR214-9 results in the loss of the compact amoeboid phenotype in the hCD8^+^ T cells. The UR214-9-treated cells align along the ICAM1 lanes, forming long, thin protrusions that stretch to up to 5-6 times the T cell amoeboid dimensions under control conditions. **b** - Zoom-in example of an UR214-9-treated hCD8⁺ T cell showing extreme cell-body elongation, centrally positioned nucleus, and longitudinal F-actin and microtubule alignment along the ICAM1 track. At full elongation, the T cell loses a morphologically distinct main body, with the nucleus (green) occupying the central region of the cell between the elongated protrusions aligned along the ICAM-1 track. *Note the extensive microtubule network caging the nucleus.* **c** - Representative tubulin images showing microtubule network expansion and enrichment after UR214-9 treatment. UR214-9-treated cells develop elongated, dense microtubule bundles extending across the full cell length and into distal protrusions, compared to compact microtubules in DMSO-treated cells. **d** - Live captures of oscillatory nuclear translocation in the UR214-9-treated hCD8^+^ T cell, spread along the ICAM1 lane (*green*). The nucleus, labeled with the live chromatin dye DRAQ5 (*magenta*), displays oscillatory displacements of up ∼55 µm within a stationary T cell boundary. See **Movie 3**. **e** - Evaluation of population-wide changes of MT network density following septin GTPase inhibition (+DMSO *vs.* +UR214-9) with live cell flow cytometry analysis of MT **(+SiR-tubulin** live dye). Inhibition of septin GTPase activity with UR214-9 enhances polymerized microtubule networks in hCD4^+^ and hCD8^+^ T cells. *Note that the MT network density increases by approximately twofold in both hCD4^+^ and hCD8^+^ T cells*. Statistical analysis: p-values were calculated using pairwise t-tests; n represents the total number of technical measurements across three biological replicates per condition.

In mesenchymal mode, cells rely on integrin-mediated adhesion ^19^ and robust traction force generation ^21–23^. Mesenchymal T cell morphology is characterized by pronounced cell elongation, arborization, and conformity to surrounding tissue architecture, as exemplified by tissue-resident memory T cells in cornea ^30^, retina ^93^, and epidermis ^4,29,94^ **(Figure 1a-2)**. These observations highlight the remarkable plasticity of T cells as they navigate the geometric confines of solid tissues to maintain tissue surveillance and residency ^95^. Consistent with this interpretation, we show that CD3-positive mouse T cells adopt an arborized morphology as they navigate through the interstitial spaces of dense mouse skin **(Figure 1c)**. Together, these observations suggest that the mesenchymal mechanotype represents a physiologically relevant adaptive strategy for T cells operating within the geometric constraints of solid tissues. However, the mechanisms that coordinate long-range force transmission along highly elongated and branched cell morphologies, thereby producing net cell displacement, remain unknown.

A third hybrid mechanical regime arises during T cell transmigration across narrow pores within epithelial or endothelial cell layers, where efficient passage requires both pushing and pulling forces to act either simultaneously or sequentially **(Figure 1a-3)**. During diapedesis, actomyosin-driven cortical compression generates hydraulic pressure that wedges the nucleus between adjacent cells, while adhesion-dependent traction forces ^96–98^ pull the nucleus through the intercellular gap, and subsequent rear cortical compression expels it onto the far side ^67,98–100^. This push-pull cooperation places transmigration at the mechanical intersection of amoeboid and mesenchymal strategies and highlights the nucleus, the largest and stiffest organelle ^27,36,61–63^, as the principal mechanical bottleneck during passage through sub-nuclear-scale constrictions **(Figure 1d)**.

**Figure 3.**
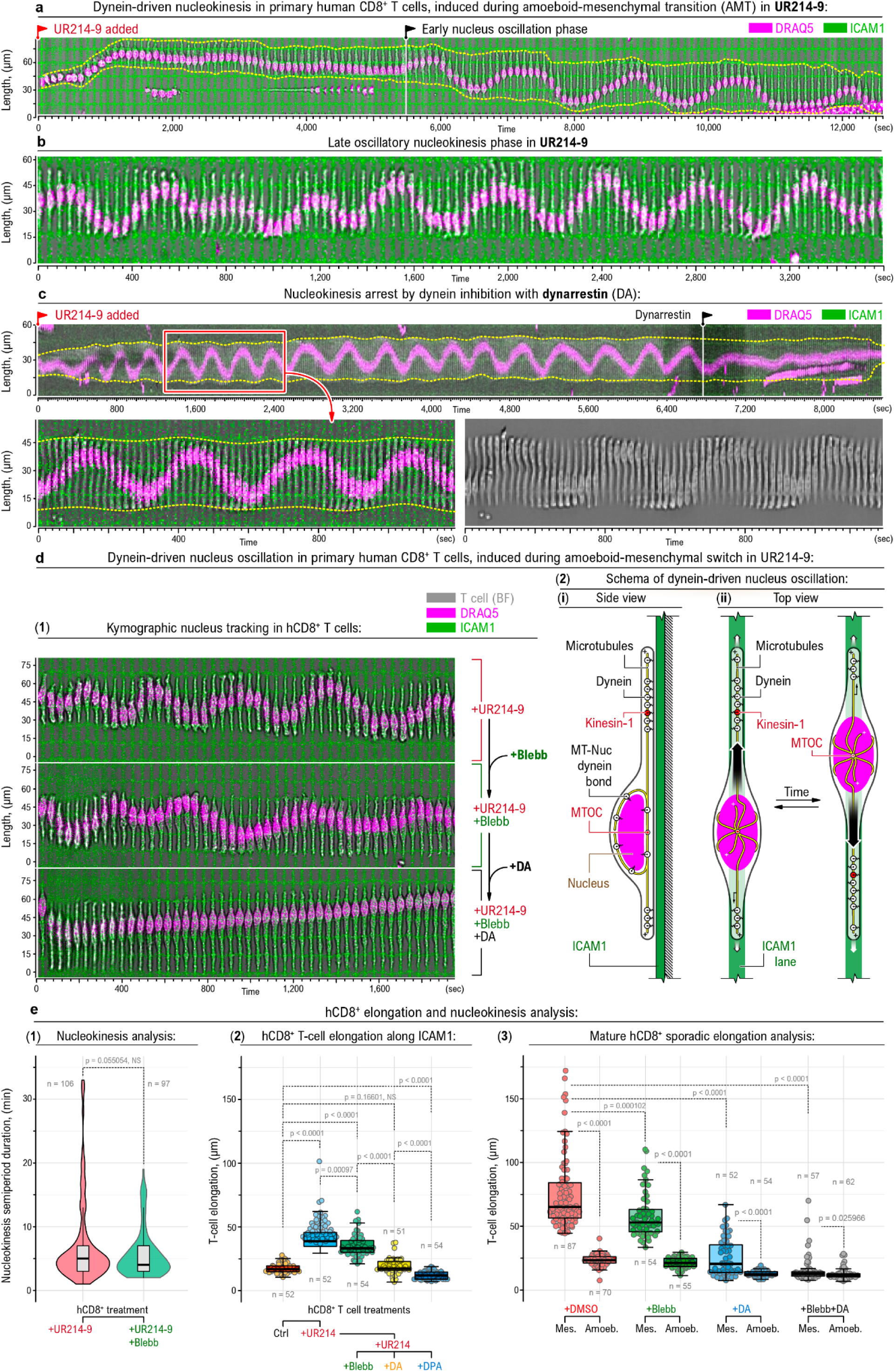
Septin inhibition induces dynein-dependent, myosin-independent oscillatory nucleokinesis decoupled from F-actin dynamics. **a** - Kymographic representation of the hCD8^+^ T cell morphological and structural dynamic transition from the amoeboid to mesenchymal phenotype upon addition of the septin GTPase inhibitor UR214-9. The treated T cell adheres and spreads its boundaries (*dashed lines*) along an ICAM1 lane within the underlying ICAM1 grid micropattern (*green*), reaching substantial elongation during the early phase of nuclear oscillation (*t* ≈ 80 min). See **Movie 4.** **b** - Kymograph of the late, stabilized phase of oscillatory nucleokinesis within the stationary boundaries of a spread T cell during septin GTPase inhibition **(+UR214-9)**. See **Movie 5.** **c** - Dynein is essential for nucleokinesis. Oscillatory nucleokinesis, induced by the UR214-9 treatment of hCD8^+^ T cells, is arrested by the dynein inhibitor Dynarrestin **(+DA)**. *Top row* - kymographic representation of stable oscillatory nucleokinesis induced by UR214-9 in a stationary hCD8^+^ T cell (cell boundary - *dashed lines*). See **Movie 6.** **d** - UR214-9-induced T cell nucleokinesis is primarily dependent on dynein and largely independent of non-muscle myosin II activity, as shown by the treatment sequence: control **(+UR214-9) →** NM myosin II inhibition **(+UR214-9+Blebb) →** dynein and NM myosin II **(+Blebb+UR214-9+DA),** See **Movie 7 and 8**. As a more stable and low-phototoxicity form of blebbistatin, we used (-)-4’ *para*-nitroblebbistatin. **(1)** Oscillatory nucleokinesis response to the sequential treatment in hCD8^+^ T cells: *top row* - stable nucleokinesis **(+UR214-9)**; *middle row* - persistent but more irregular nucleokinesis upon suppression of actomyosin contractility **(+Blebb+UR214-9)**; nucleokinesis arrest upon co-inhibition of the dynein activity **(+DA+Blebb+UR214-9)**. **(2)** Schematic representation of the nucleokinesis mechanism: the nucleus is mechanically coupled to microtubules through dynein-associated adaptor complexes, forming a nucleus-MT mechanical unit. Dynein complexes mechanically coupled to F-actin generate forces that slide MTs and translocate the attached nucleus (*nucleokinesis*). In bipolar (*i.e.*, linear, spindle-shaped) T cells, as they spread along an ICAM1 lane, dynein engagement generates nuclear motion, which induces a repolarization of actin dynamics, switching activity to the opposing protrusion, and thereby producing repeated cycles of nucleokinesis. **e** - Analysis of the hCD8^+^ T cell morphology and nucleokinesis: **(1)** Statistical analysis of the semiperiod of UR214-9-induced nucleus oscillations in septin inhibitor alone **(+UR214-9)**, and during co-suppression of the actomyosin contractility **(+Blebb+UR214-9)**. **(2)** Analysis of T cell elongation along adhesive ICAM1 lanes (*linear spreading*) in various treatment conditions, *i.e.*, control (*Ctrl*, **+DMSO),** septin inhibition **(+UR214-9)**, NM myosin II and septin inhibition **(Blebb+UR214-9)**, dynein-MT interactions and septin inhibition **(+DA+UR214-9)**, dynein motricity and septin inhibition **(+DPA+UR214-9)**. (**3**) Analysis of mature hCD8^+^ T cell spreading and elongation along the ICAM1 grids in mesenchymal (*mes*.) and amoeboid (*amoeb*.) states in control conditions **(+DMSO)**, during actomyosin contractility inhibition **(+Blebb)**, during dynein inhibition **(+DA)**, and during combined actomyosin and dynein inhibition **(+Blebb+DA)**. Statistical analysis: p-values were calculated using pairwise t-tests; n represents the total number of technical measurements across three biological replicates per condition.

Collectively, **Figure 1** establishes a conceptual framework in which the microenvironmental geometry selects for distinct T cell mechanotypes along an amoeboid-to-mesenchymal continuum. While the amoeboid program is well-suited to loosely structured environments, such as reticular 3D matrices of lymph nodes or spleen composed of discrete collagen fibers, it is mechanically insufficient for the severe confinement encountered in densely packed solid tissues and during transmigration through tight pores. We therefore focused on identifying the cytoskeletal mechanisms that enable T cells to operate beyond the limits of amoeboid motility, starting with the septin-dependent molecular switch proposed to mediate transitions between these mechanotypes ^101^.

### Septin-mediated mechanotype switch enables T cell adhesion-driven contact guidance

Similar to primary hCD4^+^ T cells ^3^, primary human CD8^+^ (hCD8^+^) T cells show a hybrid amoeboid-mesenchymal phenotype on ICAM-1 micropatterned grids, characterized by weak adhesion, limited spreading, and overall low conformity to the adhesive grid **(Figure 2a**, +DMSO). Notably, septin inhibition in hCD4^+^ T cells induces a marked mechanotype transition toward a pronounced mesenchymal mode of migration, characterized by strong integrin-mediated (LFA-1-to-ICAM-1) adhesion and spreading, extreme elongation or arborization conforming to the ICAM-1 geometry, and directional migration guided by anisotropic ICAM-1 patterns ^102^. Consequently, to investigate the mesenchymal mechanotype in hCD8^+^ T cells, we sought to suppress the amoeboid phenotype using this previously described septin-driven mechanotype switch ^102^. For this purpose, we treated activated hCD8^+^ T cells with the fluoride-stabilized form of the conventional fungal septin inhibitor forchlorfenuron (FCF) ^103^, UR214-9 ^104^. Unlike FCF, UR214-9 is more stable and exhibits lower toxicity in primary human T cells ^15^.

Treatment with 50 µM UR214-9 rapidly disrupts the hybrid state and induces a pronounced amoeboid-to-mesenchymal mechanotype transition, causing hCD8^+^ T cells to lose their compact amoeboid morphology and conform closely to the architecture of the adhesive ICAM-1 grids (**Movie 1**). This adhesion-guided elongation increases the average cell length from ∼15 µm to ∼45 µm, with individual cells reaching up to ∼90 µm, representing a three- to sixfold increase relative to the typical amoeboid dimensions and resulting in a spindle-shaped, overextended morphology **(Figure 2b**, +UR214-9). At full elongation, the cells lose a morphologically distinct central cell body and instead adopt a bipolar, spindle-like morphology, with the nucleus positioned near the center and two MT-rich protrusions extending in opposite directions along the ICAM-1 lane **(Figure 2b)**.

This septin inhibition-induced mesenchymal morphology closely resembles the reported structural features of tissue-resident memory T cells **(Figure 1a-2)** and is consistent with the arborized morphology of CD3+ T cells in mouse skin **(Figure 1c)**. Together, these observations support the idea that septin signaling functions as a molecular switch between amoeboid and mesenchymal mechanotypes.

Notably, UR214-9 treatment is accompanied by two prominent changes: (**1**) a substantial, population-wide expansion of the microtubule network (**Figures 2a-c)** and (**2**) cyclic back-and-forth nuclear translocation (live-labeled with DRAQ5, **Movie 2**) over distances reaching up to ∼60 µm, often along the entire length of spindle-shaped cells **(Figure 2d, Movie 3)**. In hCD4+ and hCD8+ lymphocytes, live-cell flow-cytometric quantification with SiR-tubulin revealed an approximately two-fold increase in MT density following septin inhibition **(Figure 2e)**. Concomitant with this increase, MTs reorganize into a cage structure surrounding the nucleus (**Figures 2b-c)**. These observations indicate that the amoeboid-to-mesenchymal transition is accompanied by extensive remodeling of the cytoskeletal architecture and suggest that the expanded MT network and associated molecular motors may contribute to the nucleokinesis ^7,105^ described below.

### Mesenchymal mechanotype transition triggers dynein-microtubule-driven oscillatory nucleokinesis

A detailed sequence of the UR214-9-induced mechanotype transition for a representative hCD8^+^ T cell on the ICAM-1 grid **(Figure 3a, Movie 4)** shows the following progression: (**1**) an initially predominantly amoeboid hCD8^+^ T cell begins adhesion-guided elongation along the ICAM-1 lane upon addition of UR214-9, followed by (**2**) a phase of nuclear oscillation (*e.g.*, at *t* = 5,500 sec).

At later times, the periodicity of the nuclear oscillation stabilizes **(Figure 3b and c, Movies 5 and 6)**, displaying regular oscillatory nucleokinesis along the stretched T cell with an average amplitude of ∼30 µm. The nucleus translocations occur within an otherwise stationary, spindle-shaped cell **(Figure 3c**, dashed line). The UR214-9-induced oscillatory nucleokinesis can persist for up to ∼2 hours during live-cell microscopy, at which point T cells experience illumination-induced phototoxicity.

The expansion of the microtubule network and its reorganization into an “enveloping nuclear cage” **(Figure 2b-c)** suggest a potential role for MT-associated motors in nucleokinesis. Consistent with this possibility, inhibition of dynein-MT interactions with dynarrestin abruptly arrests UR214-9-induced nucleokinesis but does not cause immediate cell detachment **(Figure 3c, Movie 6)**. Thus, maintenance of the elongated, adhesive morphology on ICAM1 does not require ongoing dynein activity, whereas nuclear translocation is acutely dynein-dependent. We used dynarrestin as an acute pharmacological perturbation of cytoplasmic dynein activity because it directly inhibits dynein-dependent microtubule motility, allowing rapid suppression during live-cell imaging and reducing the likelihood of compensatory cytoskeletal remodeling or secondary effects associated with longer-term genetic depletion or non-pharmacological disruption ^106^.

### Dynein drives nucleokinesis independently of actomyosin contractility

We next asked whether dynein motor activity is primarily responsible for the observed oscillatory nucleokinesis in UR214-9-treated hCD8⁺ T cells or whether conventional actomyosin contractility, driven by canonical non-muscle class-II myosin motors (NM myosin II, NMII) ^107,108^, makes a substantial contribution. To address this question, we suppressed NMII activity in T cells undergoing UR214-9-induced oscillatory nucleokinesis by treating them with the photostable, low-phototoxicity blebbistatin derivative (-)-4′-para-nitroblebbistatin during live time-lapse acquisition ^109^ **(Figure 3d-1**, +DMSO→+Blebb). In parallel endpoint/control experiments, cells were treated with (-)-blebbistatin to a final concentration of 50 µM. A 50 µM concentration of (-)-4’-para-nitroblebbistatin exceeds reported IC_50_ values for inhibition of myosin-II ATPase activity and is therefore expected to robustly suppress class-II myosin motor activity while avoiding the phototoxicity associated with conventional blebbistatin during live imaging ^110^. Since conventional actomyosin contractility in primary T cells is predominantly powered by NM myosin IIA, encoded by MYH9/NMIIA ^111^, blebbistatin-based treatment provides an appropriate strategy for suppressing NMII-dependent conventional actomyosin contractility ^112,113^.

Treatment with 50 µM (-)-4’-para-nitroblebbistatin, added to UR214-9-treated hCD8⁺ T cells during live imaging, did not abolish oscillatory nucleokinesis during the observed treatment period (>10 min; **Figure 3d-1**, +UR214-9+Blebb). Instead, nuclei continued to undergo repeated translocations between opposing protrusion tips, albeit with increased variability in the amplitude and periodicity of nuclear oscillations.

Because 50 µM (-)-4’-para-nitroblebbistatin with prolonged exposure (>10 min) is expected to robustly inhibit class-II myosin motor activity, consistent with prior live-cell evidence for rapid blebbistatin action ^114^ and biochemical measurements showing suppression of myosin-II ATPase activity ^113,115^, the persistence of nuclear translocation indicates that ongoing NMII-dependent actomyosin contractility is not the primary driver of, nor is it strictly required for, oscillatory nucleokinesis under these conditions.

Conversely, the addition of the dynein inhibitor dynarrestin (+DA) to these oscillating UR214-9- and para-nitroblebbistatin-treated cells rapidly arrested nuclear movement **(Figure 3d-1, Movie 7 and 8** +UR214-9+Blebb+DA), similar to the nucleokinesis arrest observed following dynarrestin treatment of cells exposed only to UR214-9 **(Figure 3c**, +UR214-9+DA).

Thus, dynein inhibition abolishes nucleokinesis regardless of the state of NMII-dependent actomyosin contractility. Together, these results identify dynein-generated forces as the primary driver of nuclear translocation, whereas actomyosin contractility modulates the regularity of oscillatory behavior but is not essential for its maintenance.

### Dynein slides microtubules along a stationary F-actin cortex to drive nuclear translocation

To further test the microtubule-dynein-driven nucleokinesis model, we examined the behavior of F-actin relative to the oscillating nucleus in UR214-9-treated hCD8^+^ T cells **(Supplementary Figure 1, Movie 9 and 10)**. Specifically, we tracked the positions of F-actin heterogeneities (dashed lines) within elongated protrusions and compared their dynamics with nuclear position over time.

This analysis shows that nuclear motion is decoupled from the movement of F-actin heterogeneities within the protrusions. As the nucleus moves past these F-actin heterogeneities and completes a full oscillatory cycle **(Supplementary Figure 1**, arrows), the heterogeneities continue their own slow, linear, centripetal displacement from their initial positions (arrowheads) toward the center of the elongated T cell.

If nuclear oscillation were driven primarily by actomyosin contractility, then in these highly elongated, spindle-shaped UR214-9-treated hCD8^+^ T cells, the movement of F-actin heterogeneities would be expected to remain coupled to nuclear position. Instead, the observed decoupling between F-actin dynamics and nuclear oscillation argues against a mechanism in which nuclear movement is directly driven by bulk actomyosin translocation. These observations are consistent with the involvement of a cable-like microtubule cytoskeleton and a non-myosin motor system capable of translocating the nucleus relative to the surrounding F-actin network.

A similar relationship is observed when UR214-9-treated hCD8^+^ T cells break symmetry and, instead of oscillating, begin migrating along the 1 µm-wide ICAM-1 lane. In these cells, F-actin heterogeneities remain stationary within the leading protrusions **(Supplementary Figure 2, Movie 11)**. Under these conditions, the leading protrusions display treadmilling dynamics: individual protrusion segments remain spatially stationary, while the protrusion tip extends through F-actin polymerization and the nucleus follows from the rear. These structural dynamics are likewise consistent with a model in which nuclear translocation occurs relative to a largely stationary F-actin scaffold and is driven by a microtubule-based motor system.

**Figure 3d-2** presents a working model that integrates these observations and illustrates a mechanical coupling between F-actin, dynein, microtubule, and the nucleus during oscillatory nucleokinesis, *i.e.*, cyclic nuclear movement relative to the T cell’s F-actin-defined boundaries conformed to the ICAM1 lanes. In this model, dynein motor complexes are mechanically coupled to cortical and protrusion-associated F-actin ^40,46,47,50^, which serves as a fulcrum for dynein-driven microtubule sliding along the cell cortex. As microtubules slide along the F-actin framework, the nucleus is translocated with them through its well-established mechanical linkages to the microtubule cytoskeleton ^116^. This model is motivated by the observations presented here together with prior studies demonstrating mechanical coupling between the nucleus and the microtubule cytoskeleton through dynein- and kinesin-dependent linkages ^75,105,117–119^, and showing that dynein can couple microtubules to F-actin and drive MT sliding along actin structures ^46,55^. Indeed, such coupling is now causally linked to dynein-dependent mechanical and structural coherence between microtubules and F-actin ^46,55,56,120^.

Notably, although inhibition of actomyosin contractility with 50 µM (-)-4’-para-nitroblebbistatin induces irregularity of oscillatory nucleokinesis **(Figure 3d-1**, +Blebb), it does not alter the overall periodicity of the oscillation **(Figure 3e-1**, +UR214-9→+UR214-9+Blebb), indicating that myosin activity does not solely determine the pace of nuclear oscillation. At the same time, this result suggests that actomyosin contractility contributes to the stability of the nucleokinesis cycle, possibly by stabilizing the F-actin network that serves as a mechanical substrate (fulcrum) for dynein-mediated force transmission. These observations raise the question of how dynein- and actomyosin-based contractility differentially contribute to nucleokinesis and to T cell adhesion and spreading within the microenvironment.

### Dynein and actomyosin cooperate to sustain mesenchymal T cell adhesion and spreading

Unlike the single-stride nucleokinesis of maturing neurons ^73^, mesenchymally transformed (UR214-9-treated) T cells exhibit sustained oscillatory nucleokinesis **(Figures 3a-3d)**, suggesting that dynein supports a persistent mechanical program suited to adhesion-based migration in confining environments. This is particularly relevant to T cell infiltration into solid tissues, where lymphocytes must first transmigrate through the confining boundaries of blood vessel walls during diapedesis ^121^ and then navigate tight interstitial spaces ^23,33,122^. These extreme conditions can exceed the mechanical capacity of actomyosin-driven amoeboid migration ^14,20,123,124^, overwhelming the cortical amoeboid peristalsis and nuclear propulsion mechanisms that normally sustain this mode of migration ^15,17^. These observations raise the possibility that, under such severe confinement, T cells in the mesenchymal mechanotype employ a dynamic fluid-like F-actin network to explore and conform to available space, while dynein-mediated force transmission along microtubules may contribute to nuclear propulsion and whole-cell mechanical coordination.

Consistent with this model, dynein-generated forces are known to power cancer cell migration under reduced or absent actomyosin contractility, achieved either through direct suppression of NM myosin II or through a mechanosensing-dependent failure to develop strong actomyosin contractility on compliant substrates ^40,41^. Moreover, dynein-driven cell spreading ^42^ and migration ^40^ mechanisms are particularly efficient during extensive cell elongation and branching, suitable for cell navigation under extreme confinement **(Figures 1a-2 and 1c)**. Furthermore, dysregulation of dynein complex components is associated with cancer progression, metastatic aggressiveness ^125–130^, and is used as a predictive marker of clinical outcome in multiple tumor types ^131^.

To test whether dynein contributes more broadly to the T cell mesenchymal mechano- and phenotype, we examine its role in adhesion and spreading. Dynein activity, together with actomyosin contractility, determines T cell adhesion and spreading on ICAM-1 *via* LFA-1 integrin ^132,133^. Under baseline conditions, freshly activated hCD8^+^ T cells adhere to ICAM-1 grids and adopt a compact hybrid amoeboid-mesenchymal phenotype ^3,15,102^ **(Figure 2a**, +DMSO). UR214-9 treatment disrupts this state, driving mesenchymal adhesion-guided spreading^102^ along ICAM-1 lanes and increasing average cell length from ∼18 µm to ∼40 µm **(Figures 2a-c, 3a,** +DMSO *vs.* +UR214-9) with a maximum of ∼95 µm **(Figure 2b**, +UR214-9). Pharmacological dissection of force-generating systems in UR214-9-treated cells revealed the following hierarchy: myosin II inhibition with blebbistatin produced a modest but statistically significant reduction in ICAM-1-driven cell elongation **(Figure 3e-2**, +UR214-9+Blebb), whereas dynein inhibition with either dynarrestin (DA) or dynapyrazole A (DPA) caused substantially stronger suppression of T cell spreading **(Figure 3e-2**, +UR214-9+DA and +UR214-9+DPA).

This hierarchy was confirmed in mature exhausted hCD8^+^ T cells, which can spread mesenchymally along ICAM-1 micropatterns without UR214-9 treatment: dynein inhibition, but not myosin II inhibition, strongly collapsed the elongated cell morphology **(Figure 3e-3**, +Blebb *vs.* +DA *vs.* +DPA), while only the combined inhibition of both dynein and myosin II completely abolished cell elongation **(Figure 3e-3**, +Blebb+DA). Together, these findings indicate that dynein and actomyosin make complementary but non-redundant contributions to the mesenchymal mechanotype. The dynein-microtubule system appears to provide the dominant contribution to long-range force transmission for mesenchymal spreading and nucleokinesis, whereas actomyosin supplies complementary contractile support that helps maintain the elongated morphology permissive for dynein-dependent nuclear oscillation **(Figure 3d-2)**.

### Efficient T cell transmigration requires non-redundant cooperation between dynein and actomyosin

Following characterization of the mesenchymal T cell mechanotype induced by septin GTPase inhibition and its associated dynein-driven nucleokinesis, we asked whether this mechanobiologically distinct state is better suited for migration through highly confined spaces, particularly during transmigration. To answer this question, we focused on T cell transmigration through narrow (3-µm-wide, 10-µm-long) ICAM1-coated pores, which requires substantial deformation of both the cell body and the nucleus **(Figure 4a)**, thereby exceeding the established mechanical limits of amoeboid motility ^8,134^.

**Figure 4.**
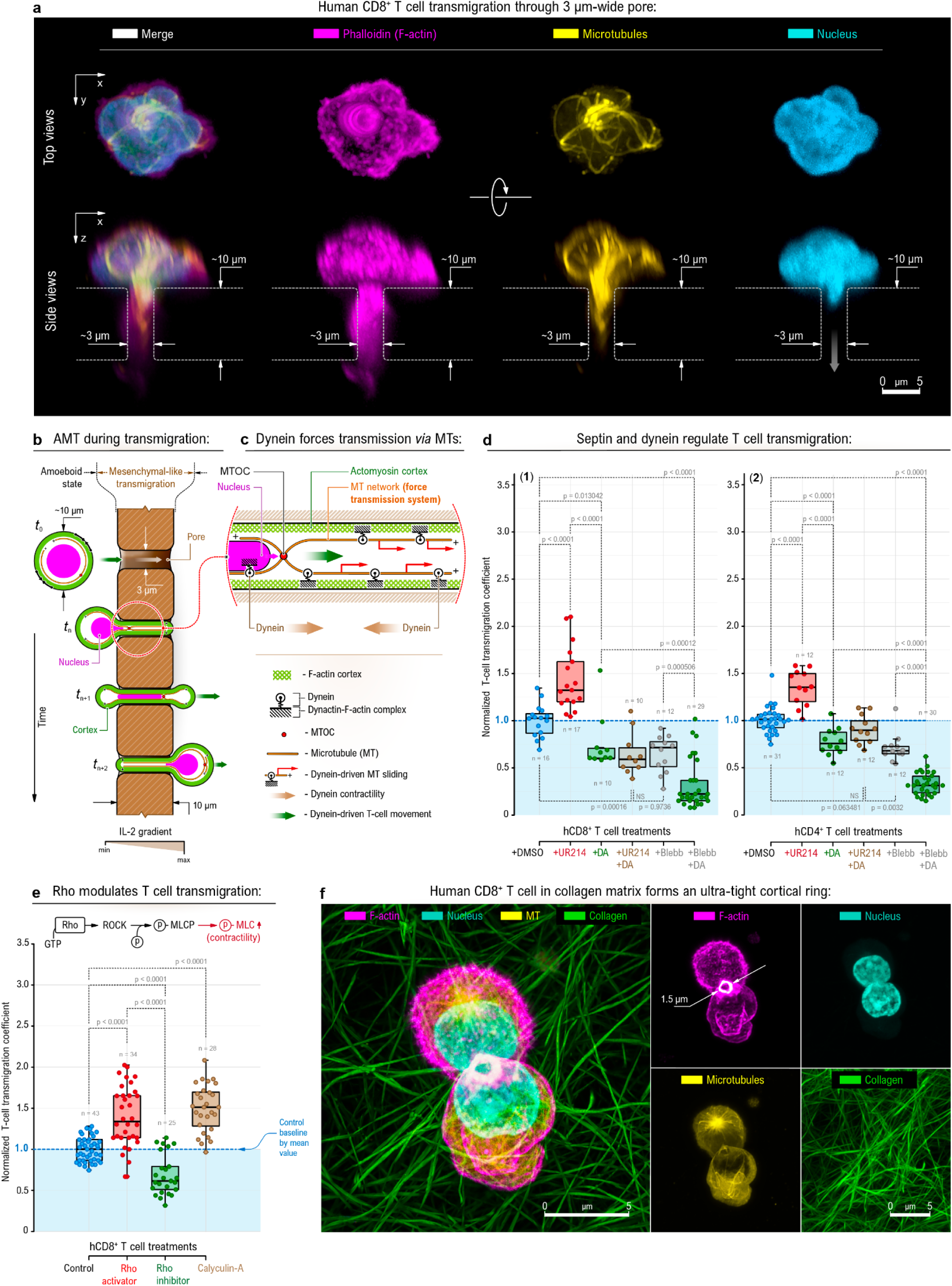
Septin inhibition enhances T cell transmigration through narrow constrictions via cooperative dynein and actomyosin force generation. **a** - Immunofluorescence 3D imaging of the human primary CD8^+^ T cell transmigrating through a 3 µm-wide, 10 µm-long pore. **b** - Schematic representation of the T cell transmigration through a 3 µm-wide, 10 µm-long pore. The pore imposes deformations on the transmigrating T cell and its nucleus that exceed those typically accommodated by the amoeboid motility mechanism. **c** - The in-pore part of the transmigrating T cell features elongation and narrowing, within which the actomyosin cortex cooperates with the microtubules: dynein complexes mechanically coupled to F-actin, *e.g.*, *via* dynactin, crosslink MTs to the actin cytoskeleton and generate pulling forces along the MT network, which acts as a system of relatively inextensible cables. These forces are transmitted to the nucleus through MT-nucleus adaptor complexes, *e.g.*, LINC. **d** - Human primary CD8^+^ and CD4^+^ T cells transmigration efficiency in response to septin inhibition **(+UR214-9)**. **e** - Enhanced hCD8^+^ T cell transmigration occurs following Rho activation or Calyculin-A treatment, both of which increase actomyosin contractility, whereas Rho inhibition decreases transmigration by suppressing myosin II activity. **f** - Bi-cameral, amoeboid (non-adhesive) human primary CD8^+^ T cell pushes its nucleus through an ultra-tight cortical ring (∼1.5 µm), demonstrating the ability of T cells to use hydraulic (amoeboid) forces to propel the nucleus through extremely narrow constrictions. Statistical analysis: p-values were calculated using pairwise t-tests; n represents the total number of technical measurements across three biological replicates per condition.

Here, we sought to test whether successful nuclear passage through a small pore requires both nucleus-pushing and -pulling forces ^100^ **(Figure 1a-3)**, with the pulling component potentially mediated by dynein-generated forces transmitted through the microtubule network **(Figures 4b, c)**. Specifically, transmigration through the tight pore may require the active contribution and mechanical cooperation of both motor systems. In this conceptual framework, dynein complexes, mechanically coupled to F-actin (*e.g.*, *via* dynactin ^46^), crosslink the microtubules to the F-actin cortex **(Figure 4c)**. This complex could function as a system of non-stretchable cables, using dynein-generated pulling forces to translocate the nucleus through narrow confinement *via* MT-nucleus adaptors, such as the LINC complex.

Notably, septin inhibition with UR214-9 significantly increased the transmigration efficiency of both primary hCD8^+^ and hCD4^+^ T cells by ∼30% **(Figure 4d**, +UR214-9), indicating that this mechanotype switch primes T cells for navigation through highly confined spaces. This enhancement is consistent with the dynein-dependent nucleokinesis induced by septin inhibition **(Figure 2c)**, and coordinated force transmission through the microtubule cytoskeleton.

To further investigate the force-generating mechanisms governing T cell transmigration, we examined the individual and combined contributions of actomyosin and dynein activity. Individual inhibition experiments revealed that dynein and actomyosin make cooperative, non-redundant contributions to transmigration **(Figure 4d)**. Selective inhibition of either dynein (+Dynarrestin, DA) or NM myosin II ((-)-4’-para-nitroblebbistatin) alone only partially suppressed transmigration by 40-45% in both primary hCD4^+^ and hCD8^+^ T cells. In contrast, combined dynein and myosin inhibition (+Blebb+DA) produced the strongest suppression, approaching nearly complete abrogation of transmigration **(Figure 4d**, +Blebb+DA).

Notably, dynein inhibition with dynarrestin fully abolished the UR214-9-induced enhancement of transmigration in hCD4^+^ and hCD8^+^ T cells **(Figure 4d**, +DA+UR214-9). Specifically, transmigration levels fall below the untreated control baseline **(Figure 4d**, +DMSO) and approach the levels observed with dynarrestin treatment alone **(Figure 4d**, +DA), indicating that dynein activity is required for the UR214-9-enhanced T cell transmigration.

Together, the results indicate that both pushing and pulling forces contribute critically to efficient transmigration. In the model outlined in **Figure 1a-3**, actomyosin contractility and dynein-generated forces, relayed *via* microtubule cables, cooperate during transmigration: the actomyosin cortex hydraulically pushes the nucleus into the pore ^17^, while the dynein-microtubule system contributes a pulling component. This interpretation is consistent with our finding that enhanced actomyosin contractility, achieved either by RhoA activation (+Rho activator II) or Calyculin A, substantially increases hCD8^+^ T cell transmigration, whereas suppression of RhoA (+Rho inhibitor I) or myosin II ((-)-4’-para-nitroblebbistatin) partially reduces transmigration **(Figure 4e)**. The relevance of hydraulic pushing forces is further supported by the ability of T cells to use such forces to push their nucleus through ultra-tight cortical rings (∼1.5 µm) **(Figure 4f)**. Together, these findings indicate that efficient T cell transmigration through highly confined spaces requires mechanical cooperation between actomyosin contractility and dynein-microtubule-based force transmission.

### Integrative mechanochemical model recapitulates self-organized nuclear oscillations

We used computer simulations to test whether a minimal mechanochemical model coupling cell and nuclear shape dynamics, cell polarization, and dynein-microtubule (MT) forces could reproduce the stable oscillatory nucleokinesis observed experimentally **(Figure 5a-1)**. To that end, we expanded our previously developed cellular morphodynamics framework ^135,136^, which represents the cell as a finely pixelated domain whose boundary evolves through stochastic protrusion-retraction events regulated by actin polymerization (reaction-diffusion module), focal adhesions (agent-based module), curvature, and cell area, as described in detail in **Methods**.

**Figure 5.**
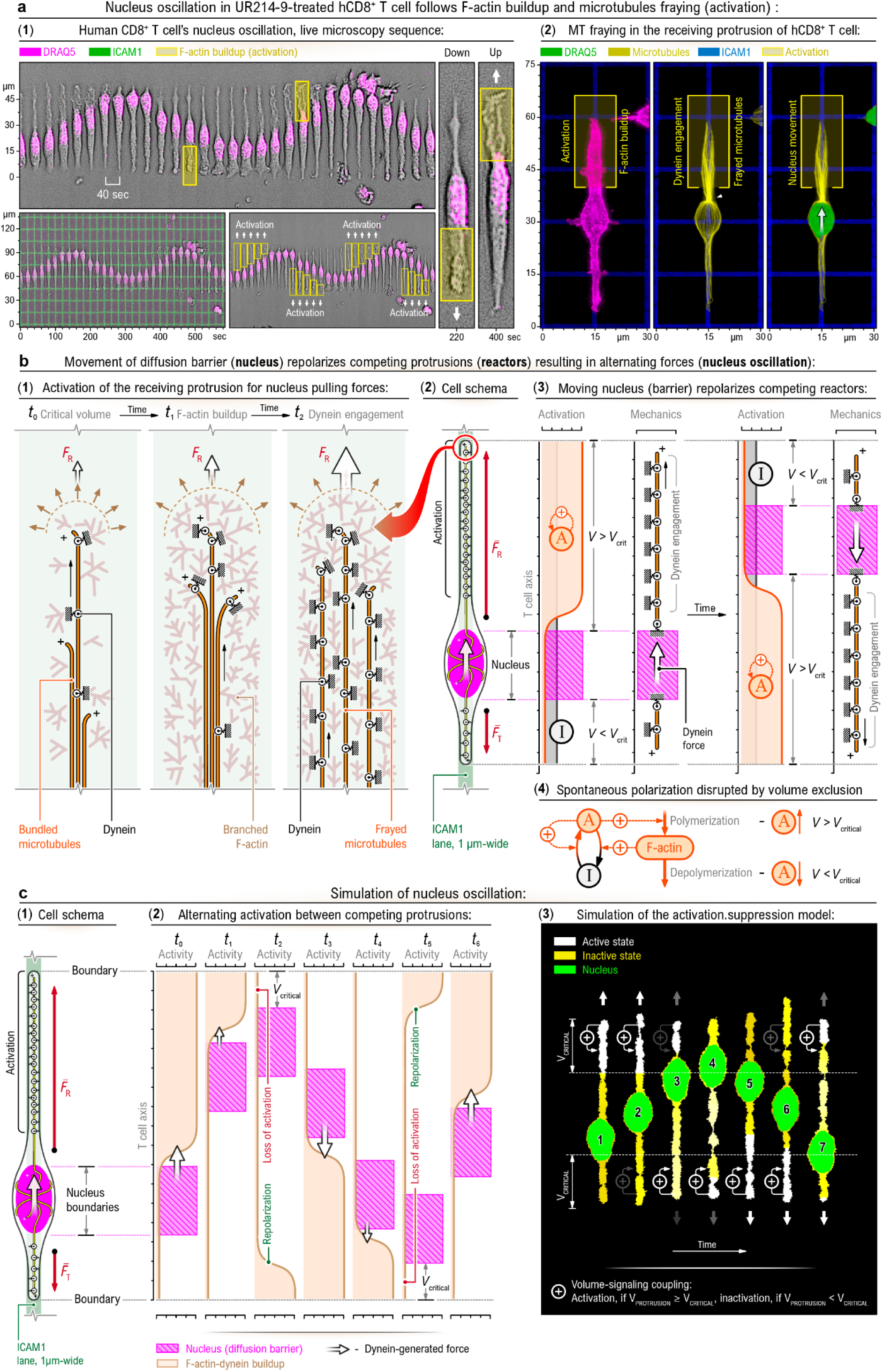
Spontaneous polarization and nucleus-mediated volume exclusion drive alternating dynein engagement and nuclear oscillations. **a** - Live-cell imaging and structural correlates of nuclear oscillation (See **Movie 12**): **(1)** Time-lapse live microscopy sequence of a UR214-9-treated human CD8^+^ T cell adhered, linearly spread, and elongated along a 1-µm-wide ICAM1 lane shows repeated oscillatory movement of the nucleus (DRAQ5, *magenta*) between two opposing protrusions. Selected boxed regions (*yellow*) highlight transient F-actin accumulation events associated with nuclear displacement before (down) and after (up) directional reversal of nuclear motion. The associated kymographs below delineate the spatial arrangement of the adhesive ICAM-1 micropattern (*bottom left*) and alternating activation dynamics (*boxed regions*) between the protrusions (*bottom right)*. **(2)** High-resolution immunofluorescence (*3D-stack, maximum projections*) displays receiving protrusion structural changes, associated with protrusion activation: F-actin buildup within the receiving protrusion is associated with increased MT-F-actin interpenetration (fraying) throughout the protrusion volume and precedes nuclear advance into the receiving compartment. **b** - Mechanistic model for nucleus-driven alternation between competing protrusions. **(1)** Sequential model of activated actin polymerization and dynein engagement. At *t*_0_, the protrusion is below a critical volume and contains mostly bundled microtubules with limited force-producing capacity. At *t*_1_, F-actin accumulation and protrusion expansion increase the extent of MT-F-actin interpenetration, resulting in microtubules “fraying” within the enlarged protrusion volume (*t*_2_). F-actin-dynein complex-generated pulling forces in the receiving protrusion (*F*_R_) drive nuclear translocation. **(2)** Cell schema summarizing force balance during T cell’s nuclear oscillation on a 1-µm ICAM1 lane. The actin-rich(frayed) receiving protrusion with engaged dynein generates a dominant pulling force (*F*_R_), whereas the opposite protrusion exerts a weaker trailing force (*F*_T_). The nucleus acts as a central diffusion and mechanical barrier between competing protrusions; consequently, nuclear position influences the effective extent of both protrusions. **(3 and 4)** Conceptual *activation-mechanics* model showing how movement of the nucleus repolarizes protrusion activity. When a protrusion exceeds the critical activation volume (*V* > *V*_crit_), the F-actin polymerization-regulating signal “A” forms spontaneous excitation and initiates F-actin buildup and dynein engagement, which enables the development of the nucleus-pulling force. As the nucleus advances into such an activated (receiving) protrusion, it reduces the available protrusion volume and suppresses activation signal “A” into the inactive state “I”, while increasing the effective volume of the opposite protrusion, thereby enabling repolarization and force reversal. **(4)** Minimal signaling circuit for spontaneous polarization of an activator of F-actin polymerization and its disruption by volume exclusion. Local protrusion volume *V* above the critical value (*V* > *V*_crit_) maintains the upstream F-actin polymerization regulator (represented generically as signal “A”) in the active state. Nuclear encroachment reduces the available volume below the critical threshold (*V* < *V*crit), promoting its inactivation (“I” state), the depolymerization of F-actin, and resetting the polarization process. **c** - Simulation of nuclear oscillation based on activation-mechanics coupling. **(1)** Cell geometry emerged in simulation due to the constraining of adhesion dynamics to a 1-µm ICAM1 lane, resulting in a spindle-shaped T cell with the nucleus compartmentalizing its volume into the opposing protrusions. **(2)** The emerging sequence of alternating dynein engagement between two competing protrusions. Nuclear position, activation of F-actin polymerization, and dynein-generated force change cyclically as nuclear displacement suppresses polarization patch in the currently occupied protrusion and permits reactivation of the opposite side. **(3)** Simulation output of the *activation-mechanics* model showing oscillatory nucleokinesis between alternating activation sites of an F-actin regulator in opposed protrusions. Green, nucleus; white, active protrusion state; yellow, inactive protrusion state..

We introduced two key extensions. First, the nucleus was modeled explicitly as an inner deformable domain with defined rigidity. Second, dynein-MT-F-actin interactions were represented as an effective pulling field acting on the nucleus and spatially coupled to regions of elevated polarity activity. Without explicitly imposing oscillatory behavior, the model reproduced the experimentally observed sequence in which spontaneous polarization drives F-actin accumulation **(Figure 5a-2, Movie 12)**, dynein-mediated pulling translocates the nucleus toward the active site **(Figure 5b-1,2)**, and nuclear arrival triggers local repolarization through cytoplasmic volume exclusion, reducing activity at that site **(Figure 5b-3)**. This minimal coupling between spontaneous polarity, dynein-mediated pulling, and nucleus-triggered depolarization was sufficient, without introducing additional regulatory assumptions, to generate sustained, self-organized nuclear oscillations between opposing protrusions **(Figure 5c)**. The resulting dynamics were robust and closely resembled the experimentally observed rhythmic nucleokinesis **(Movies 13)**. Importantly, oscillatory nucleokinesis was not imposed but emerged from the coupling between spontaneous polarity dynamics, dynein-mediated nuclear translocation, and nucleus-induced local suppression of actin polymerization. Mechanistically, oscillations emerged from a feedback loop in which the arrival of the nucleus disrupts the local polarity site, while the excitable reaction-diffusion system establishes a new polarity patch at a distant location (typically at the tip of an opposing protrusion with high curvature and sufficient volume: *V* > *V*_crit_), thereby reversing the direction of nuclear motion.

Finally, the qualitative pattern of nuclear oscillations depended on how dynein engagement was coupled to the filamentous actin cytoskeleton and polarization dynamics. Alternative assumptions regarding the timing and spatial distribution of dynein-mediated forces (see Methods) produced distinct oscillatory behaviors, including symmetric motion and asymmetric acceleration or deceleration phases **(Figure 6a and 6b, Movie 14 and 15)**. These simulated patterns recapitulated the range of oscillatory behaviors observed experimentally **(Figure 6c)**, indicating that differences in dynein engagement and force transmission are sufficient to generate the diversity of oscillatory behaviors.

**Figure 6.**
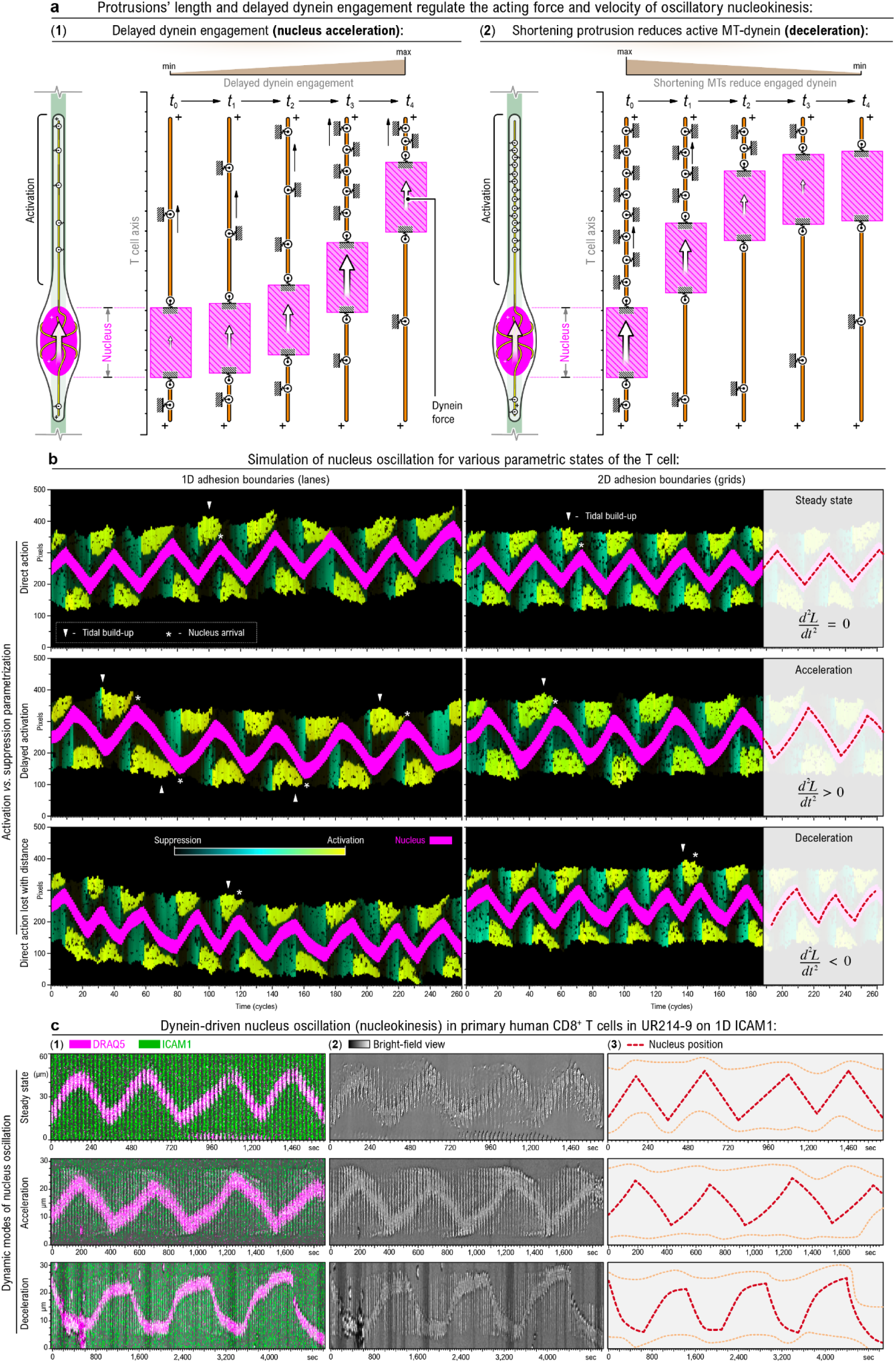
Protrusion length and delayed dynein engagement regulate the kinematics of oscillatory nucleokinesis. **a** - Mechanical models of nuclear acceleration and deceleration. **(1)** Schematic of delayed dynein engagement in the activated (frayed) protrusion leading to nucleus acceleration. As the receiving protrusion forms a high-activity site of actin polymerization, dynein recruitment to the actin cortex and engagement with MTs are initiated but take time to develop before the pulling force begins to generate significant nuclear displacement. Therefore, the progressive engagement of dyneins over time (***t*_0_** through ***t*_4_**) generates an increasing cumulative pulling force ***F*_R_**, resulting in acceleration of nuclear velocity. **(2)** Schematic illustrating how shortening of the receiving protrusion causes nuclear deceleration. As the nucleus approaches the distal end of the cell’s receiving protrusion, the available MT “track” length decreases (***t*_0_** through ***t*_4_**). The reduction in MT length limits the number of engaged dynein motors, leading to a reduction in pulling force and a characteristic slowing (deceleration) of the nucleus as it reaches the protrusion tip. **b** - Kymographs of nuclear oscillations generated by the computational model (Figure 5c**(3)**; **Methods)**. The activation level of the regulator of actin polymerization is shown using a black-to-cyan-to-yellow (low-to-high) colormap. Three rows illustrate three different assumptions regarding dynein engagement: *Top row* - The direct activation model: the dynein motor complexes are engaged immediately with the activation of the actin polymerization component. See **Movie 13.** *Middle row* - The delayed activation model: the component representing dynein engagement accumulates over time at the area of high activity of the actin polymerization component. See **Movie 14.** *Bottom row* - The direct activation with the length-dependence model accounts for dynein motors engaged along protrusions, with the resulting MT-pulling force proportional to protrusion length. See **Movie 15.** The nucleus is shown in magenta, and its motion pattern depends on the assumptions regarding dynein engagement. In the “delayed activation” case (*top row*), upon the polarization switch, the nucleus begins to move slowly toward the newly formed polarity patch in the receiving protrusion, and its speed increases as the dyneins reach full engagement (“acceleration” pattern in the top row). In the “direct activation” case (*middle row*), the nucleus moves with a nearly constant speed between the protrusion tips every time polarity switches (“constant velocity” pattern in the middle row). In the “direct activation with length dependence” (*bottom row*), the nucleus slows down as it approaches the protrusion tip because fewer engaged dynein motors remain within the shortening protrusion (“deceleration” pattern in the bottom row). Two types of ICAM1 substrates (grids) were considered: vertical parallel ICAM1 lanes (1D, *left column*) and crisscrossed vertical and horizontal ICAM1 lanes (2D, *right column*). In the simulation, the crossed ICAM1 lanes are assumed to promote more stable cell attachment at the intersection points. As a result, the overall cell shape drifts up or down during nucleus oscillations on the 1D ICAM1 lanes. In contrast, cell protrusions are mostly stalled on the 2D grids at the intersections with the horizontal strips, while the nucleus continues to oscillate between these stabilized protrusion anchoring points. Both types of simulations are consistent with the corresponding 1D and 2D experimental setups. **c** -Experimental kymographs reveal multiple patterns of oscillatory nucleokinesis in the human primary CD8^+^ T cell along the 1D ICAM1 1µm-wide lane that qualitatively correspond to the alternative model assumptions shown in panel b. **(1)** - Fluorescence imaging of the oscillating nucleus (*magenta*, DRAQ5-labeled) and the underlying adhesive 1µm-wide ICAM1 lane (*green*). **(2)** Brightfield kymographs, **(3)** Kymographs outlines of nuclear movement and cell boundaries (*dashed lines*). *Top row* - Zig-zag nuclear motion with approximately constant velocity between turning points, resembling the direct activation model. See **Movie 13**. *Middle row* - Accelerating nuclear motion, in which velocity increases as the nucleus approaches the turning point, resembling the delayed activation model. See **Movie 14**. *Bottom row* - Decelerating nuclear motion, in which velocity decreases as the nucleus approaches the turning point, resembling the direct activation with length-dependence model. See **Movie 15.**

### Oscillatory nuclear translocation underlies stride-wise saltatory migration in mesenchymal T cells

Oscillatory nucleokinesis, once established within the elongated T cell body, does not merely represent repeated back-and-forth nuclear displacement. Instead, it functions as the fundamental kinematic unit underlying net cell translocation in mesenchymally transformed hCD8⁺ T cells. Time-lapse imaging of UR214-9-treated hCD8⁺ T cells migrating on ICAM-1-patterned substrates reveals that cells advance through a characteristic saltatory, stride-wise pattern of locomotion **(Figure 7a, Movie 16)**, meaning that rather than undergoing smooth, continuous translocation, cells progress through discrete migratory events, each anchored by a nuclear displacement between competing dynamic protrusions.

**Figure 7.**
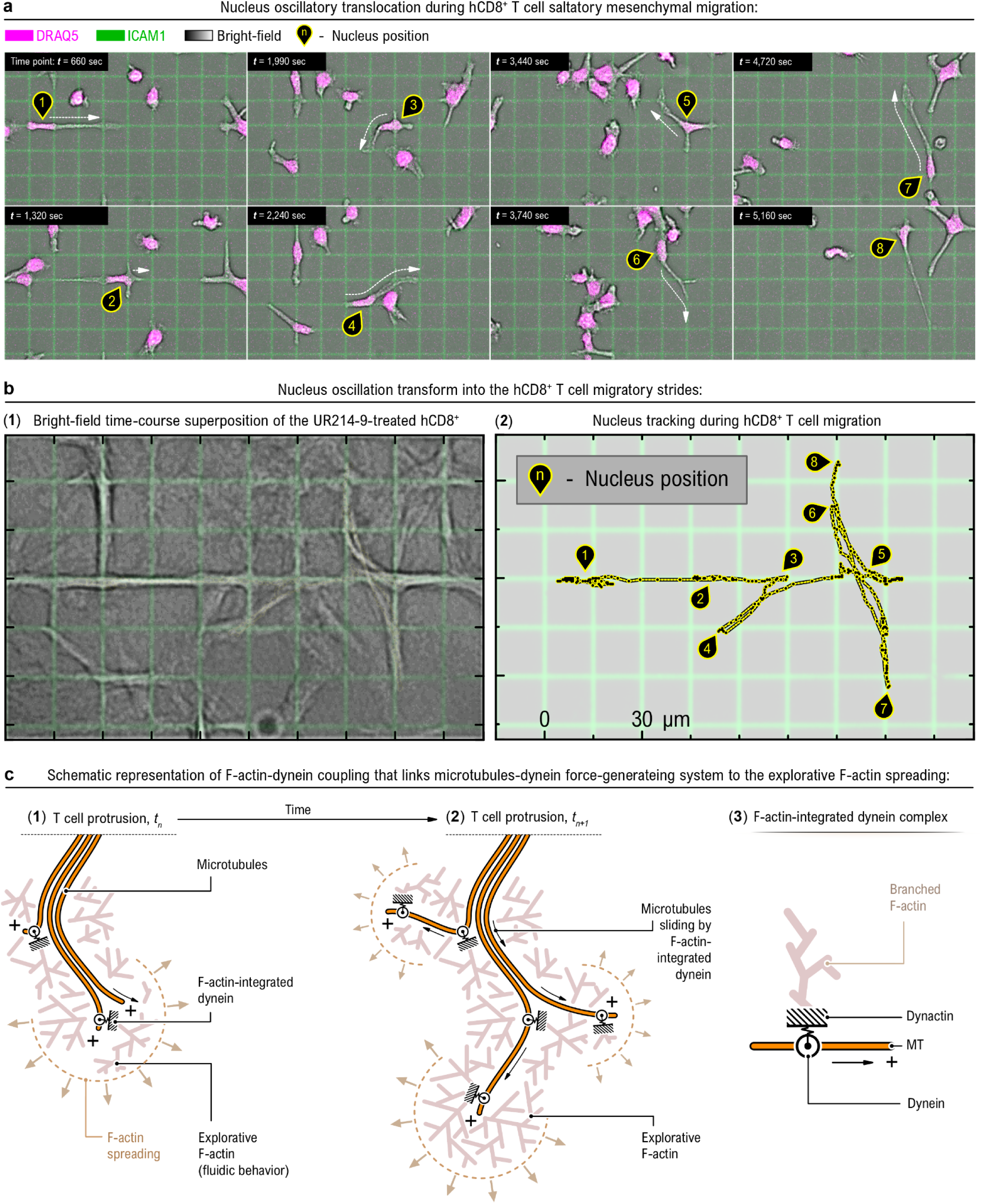
Oscillatory nuclear translocation underlies saltatory mesenchymal migration in primary hCD8⁺ T cells. **a** - Time-lapse of mesenchymal T cell migration (See **Movie 16**). Fluorescence and bright-field imaging of UR214-9-treated hCD8⁺ T cells migrating on an ICAM1-patterned substrate (ICAM1, *green*; chromatin/DRAQ5, *magenta*; bright-field, *gray*). Numbered markers (**1-8**) and white arrows track successive nuclear positions across eight time points (*t* = 660-5,160 sec), revealing a characteristic saltatory (stride-wise) mesenchymal migration pattern. The nucleus undergoes repeated oscillatory translocation between competing protrusions as the cell alternately explores and advances along the adhesive grid. At each repositioning step, the trailing protrusion undergoes a slingshot-like detachment, releasing tension and enabling the nucleus to stride forward into the nascent leading protrusion. **b** - Nuclear oscillations resolve into net forward displacement during saltatory migration. (**1**) Bright-field time-course superposition of UR214-9-treated hCD8⁺ T cells, illustrating the sequence of exploratory protrusions that collectively define a continuous migratory track. (1) Single-cell nucleus tracking (n, pin marker) maps the oscillatory stride trajectory over the same imaging period, with numbered waypoints (**1-8**) corresponding to successive nuclear positions. The irregular, back-and-forth oscillation pattern results in net directional displacement, consistent with stride-wise nuclear translocation serving as the elementary unit of saltatory cell movement. **c** - Mechanistic model linking the microtubule-dynein force-generating system to exploratory F-actin spreading. (**1**) At time ***t*□**, nascent protrusion formation is supported by exploratory F-actin spreading (fluid-like behavior at the advancing front) and F-actin-integrated dynein complexes that engage microtubules emanating from the cell body. (**2**) At time ***t***□₊₁, F-actin-integrated dynein drives microtubule sliding within the protrusion, generating forces that advance the protrusion tip, displace the nucleus, and contribute to net cell movement. Continuous exploratory F-actin spreading at the front sustains protrusion elongation throughout this process. (**3**) A possible elementary molecular ensemble of an F-actin-integrated dynein complex. Branched exploratory F-actin recruits dynactin, which activates dynein on the microtubule (MT) lattice and directs force toward the MT plus end (+), mechanically coupling the actin and microtubule systems to coordinate nuclear translocation with protrusion-driven cell migration.

The nucleus, driven by dynein-generated forces transmitted through the microtubule network, translocates forward into a nascent leading protrusion, followed by slingshot-like rear-protrusion detachment in which rear adhesion is released and the cell body repositions around the newly advanced nucleus. This stride-wise behavior is particularly pronounced in mature, exhausted hCD8⁺ T cells, which display a higher frequency of rear-protrusion adhesion failures, consistent with a lower threshold for rear detachment. Nucleus tracking across individual cells confirms that locally bidirectional oscillations resolve into net forward displacement over time **(Figure 7b)**, with each stride contributing a net directional increment along the adhesive guidance cue.

Analysis of the underlying cytoskeletal structural dynamics provides mechanistic insight into how each stride is generated **(Figure 7c)**. At stride onset (*t_n_*), the leading protrusion’s exploratory dynamics are supported by fluid-like dynamic F-actin network spreading at its advancing front, while F-actin-integrated dynein complexes engage microtubules emanating from the perinuclear region, establishing the mechanical linkage required for force transmission. As the stride matures (*t_{n+1}_*), the nucleus translocates toward the leading protrusion, consistent with dynein-mediated force transmission through the microtubule network, while continuous F-actin polymerization at the protrusion tip sustains elongation. At the molecular level, these dynamics are consistent with previously described interactions in which branched F-actin recruits dynactin, thereby promoting dynein activation and mechanically coupling the actin and microtubule cytoskeletons. Together, these observations establish oscillatory nuclear translocation as a defining kinematic signature of mesenchymal migration in this T cell state and support a model in which the dynein-microtubule-F-actin axis serves as the primary mechanical engine driving saltatory locomotion.

### Dynein-dependent protrusion dynamics are observed in living tissue

Our current findings establish dynein as a primary driver of nucleokinesis and transmigration in primary human T cells operating under a mesenchymal mechanotype. To assess whether dynein-dependent force generation extends more broadly to other tissue-resident immune cells, we examined dynein’s role in regulating protrusion dynamics in Langerhans cells. Langerhans cells are particularly well-suited for this analysis because they reside within the epidermis, a mechanically confining, tightly packed epithelial environment **(Figures 8a, b)**. Langerhans cells continuously extend and retract dendritic protrusions to survey the surrounding tissue for antigens ^31,137,138^. This persistent protrusion remodeling shares functional similarities with the exploratory protrusive behavior observed in mesenchymally transformed T cells **(Figures 5 and 7)**, suggesting that a shared dynein-dependent cytoskeletal program may underlie protrusion-driven motility across distinct tissue-resident immune cell types.

**Figure 8.**
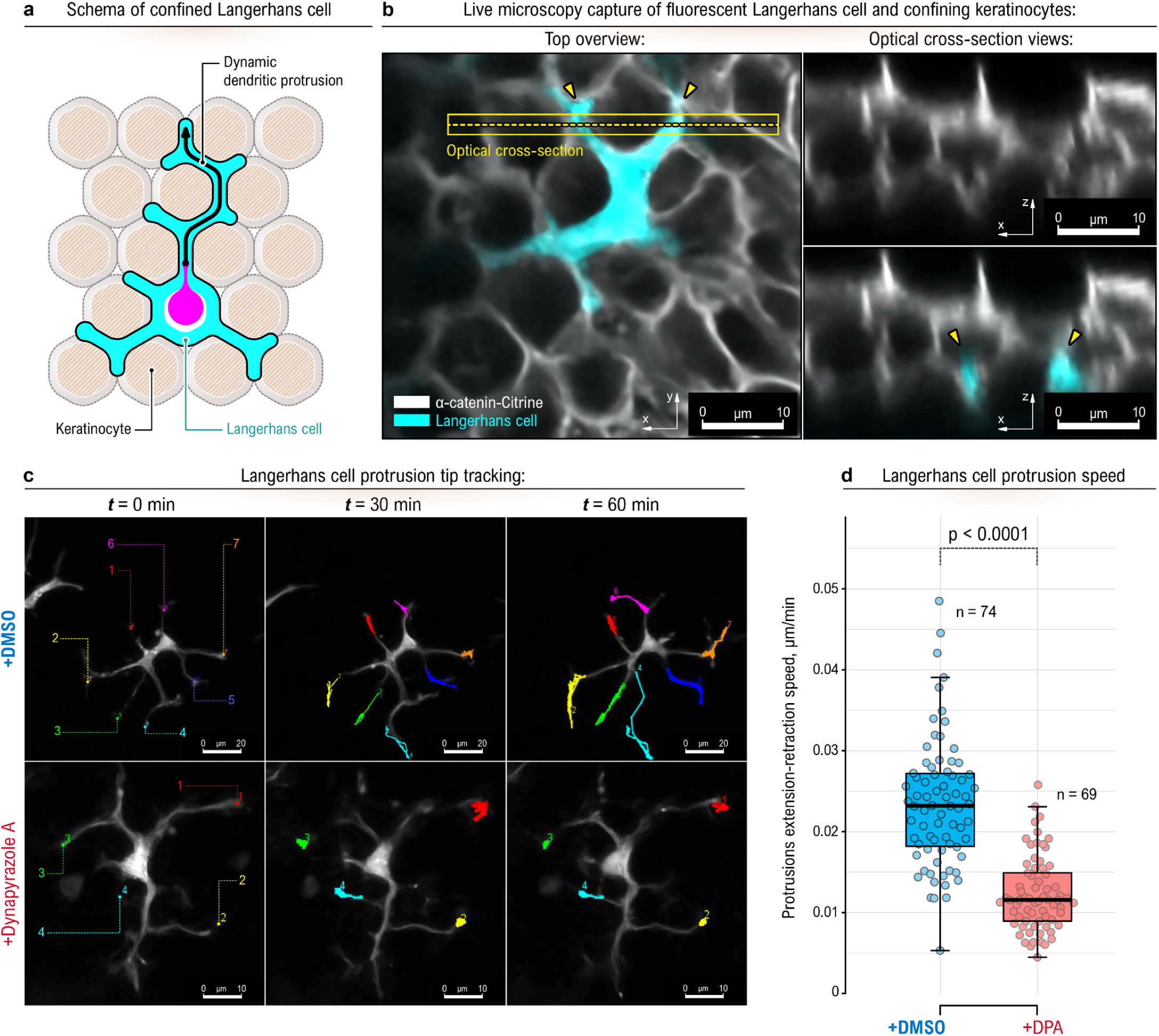
Dynein inhibition suppresses protrusion dynamics in tissue-resident Langerhans cells in situ. **a** - Schematic of a Langerhans cell extending dendritic protrusions throughout the confined epidermis composed of adherent keratinocytes. **b** - Representative confocal micrograph of a Langerhans cell (labeled with *Tg(mpeg1.1:YFP)*) extending protrusions (arrowheads) between basal cell membranes (labeled with *Gt(ctnna-citrine)*) of the zebrafish epidermis. **c** - Quantification of Langerhans cell protrusion tip extension-retraction speed under control conditions (+DMSO) and following dynein inhibition with Dynapyrazole A (+DPA), see **Movie 17**. Each dot represents a tracked protrusion tip; box plots show the median and interquartile range, with whiskers indicating the spread of the distribution. Dynein inhibition significantly reduces protrusion tip dynamics relative to control (**p** < 0.0001; +DMSO, *n* = 74 protrusions; +DPA, *n* = 69 protrusions). **d** - Representative time-lapse images of Langerhans cells with color-coded tracking of individual protrusion tips over a 60-min imaging period under control conditions (+DMSO, top) and after treatment with Dynapyrazole A (+DPA, bottom). Colored dashed lines at *t* = 0 min indicate the initial positions of tracked protrusion tips, and colored trajectories at later time points show their subsequent displacement. Under control conditions, protrusion tips undergo pronounced extension-retraction dynamics, whereas dynein inhibition markedly restricts protrusion tip motility and overall protrusion remodeling. Scale bars, 20 µm (top) and 10 µm (bottom).

Skin explants from adult zebrafish provide an experimentally accessible system for examining the effects of cytoskeletal perturbations on Langerhans cell dynamics within the epidermis ^31,139,140^. To test if Langerhans cell protrusion motility is dynein-dependent, we treated zebrafish skin explants with the dynein inhibitor Dynapyrazole A and quantified protrusion dynamics via time-lapse microscopy **(Figures 8c, d, Movie 17)**. Quantification of protrusion-tip displacement velocities across a population of tracked protrusions (n = 74) confirmed robust baseline protrusion dynamics **(Figure 8c**, +DMSO). Specifically, under control conditions (+DMSO), Langerhans cell protrusion tips exhibited pronounced and rapid extension-retraction cycles over a 60-minute imaging window, consistent with active cytoskeletal remodeling and dynamic protrusion turnover **(Figure 8d**, top).

Dynein inhibition with DPA produced a marked and statistically significant reduction in protrusion tip motility relative to control (p < 0.0001; n = 69 protrusions; **Figure 8c**, +DPA). Time-lapse imaging corroborated this finding, revealing that DPA-treated Langerhans cells exhibited substantially restricted tip displacement of protrusions and diminished overall capacity for protrusion remodeling compared to controls **(Figure 8d**, bottom), reinforcing the hypothesis that dynein activity broadly contributes to exploratory motility in immune cells residing within mechanically confining tissue environments.

## DISCUSSION

Pioneering research by Jurii Vasiliev and Yuri Gelfand established that microtubules are essential for mesenchymal motility ^141^. Our study revisits the original ideas and extends the conceptual framework of mesenchymal cell motility ^142^ into a regime of low actomyosin contractility ^40,143–145^, which is required to achieve pronounced morphological compliance ^39,40,144,146,147^ and to enable cells to navigate within spatially complex and often severely confining environments ^143–145^. These conditions are directly relevant to the persistent challenge of T cell infiltration into solid tumors ^81–85,122^, where cytotoxic T cells must squeeze through interstitial spaces ^1,29,30^ that are often too narrow to traverse ^8,15,134^. Our study establishes the dynein-microtubule (MT) force-transmission axis as a fundamental mechanobiological mechanism driving nuclear translocation during confined T cell migration. We demonstrate that this axis functions independently of canonical actomyosin contractility to navigate physical environments that exceed the limits of amoeboid motility. By integrating acute pharmacological dissection with computational modeling, we show that dynein-dependent nucleokinesis is a self-organized oscillatory process critical for cell navigation.

The demand for morphological adaptiveness, while striking in the immune context, is in fact shared across many fundamental biological processes. Neural crest cells navigate tightly packed embryonic tissue ^148^, neuron growth cones ^58^, and dendrites extend through the crowded environment of the nervous system ^149,150^, and metastatic cancer cells infiltrate dense healthy tissues ^13,40,151^. In particular, for cancer cells, migration within dense 3D collagen depends on dynein and microtubules ^42^. Clinical studies also repeatedly associate dysregulation of microtubule motors and their cofactors with cancer metastasis ^125–130^. Bioinformatic analysis ^152^ and genomic profiling ^131^ link elevated expression of dynactin’s subunit-5 and dynein’s heavy chain-1 with a substantially lower survival rate for breast cancer patients. Upregulation of dynactin’s subunit-2 (DCTN2) and/or subunit-5 (DCTN5) positively correlates with the metastatic aggressiveness of breast cancer ^153^. Pharmacological disruption of mechanical forces generated by dynein ^50,154–159^ and kinesin ^74,160,161^ impairs MT-dependent breast cancer cell shape and contact guidance ^37,38^. Collectively, these findings strongly implicate microtubules and their motors in confined cell migration.

Using primary human cytotoxic T cells as our experimental model, we show that the F-actin-linked dynein-microtubule force-generating machinery is an essential yet previously overlooked mechanobiological program that enables nuclear propulsion during confined migration and transmigration. This program emerges when the canonical amoeboid mechanotype is suppressed or no longer effective. Pharmacological inhibition of septins, key structural and signaling regulators of the amoeboid mechanotype ^40,89^, with the UR214-9 inhibitor ^104^, acts as a molecular switch, not only disrupting the amoeboid program ^15^ but also initiating a distinct mesenchymal mechanotype, enhancing adhesion-driven cellular elongation and triggering dynein-powered oscillatory nucleokinesis through an expanded microtubule network **(Figure 2c)**.

Dynein’s principal role in T cell locomotion and nucleokinesis is established by converging orthogonal evidence: suppression of dynein motricity with dynarrestin halts oscillatory nucleokinesis regardless of non-muscle myosin II’s activity status, indicating that dynein-generated forces alone are sufficient for nuclear displacement. On the other hand, mesenchymal-like T cells’ structural dynamics, in which F-actin flows centripetally within the T cell protrusions while the nucleus rapidly oscillates independently of F-actin flow, indicate that T cells may utilize cable-like microtubules for long-range transmission of dynein-generated forces, coupled to F-actin *via* cytoskeletal adaptors like dynactin. While mechanistically similar to dynein-driven nuclear movement in neural precursors and other developmental contexts ^74^, T cell nucleus motility is repetitive, suggesting an exploratory function of nucleokinesis in tissue navigation.

Our dissection of nucleokinesis relied on a panel of acute pharmacological perturbations rather than a genetic approach, a choice motivated by the need to isolate the immediate, motor-specific contribution of each cytoskeletal system to nuclear translocation. Genetic depletion strategies, shRNA, CRISPR-based knockout, or inducible degron systems, typically require hours to days to achieve effective protein loss, a timescale over which primary T cells undergo perturbation-associated phenotypic drift with compensatory cytoskeletal remodeling that obscures the specific motor contribution under study. Acute pharmacology circumvents both problems: each cell can serve as its own before/after control within a single imaging session, and the short window between drug addition and phenotypic readout limits the effects of secondary, compensatory reorganization of the cytoskeleton.

Within this pharmacological framework, each compound was selected to target a mechanistically distinct node of the cytoskeletal machinery implicated in nucleokinesis. To perturb septin-dependent cortical organization, we used UR214-9, a chlorinated analog developed through structure-activity relationship optimization of the parent compound, forchlorfenuron (FCF), which inhibits proliferation in the low single-digit micromolar range ^104^. Prior work from our own group established that acute inhibition of septin GTPase activity with this compound is sufficient to rapidly displace septins from the cortical actomyosin, consistent with an immediate disruption of septin-actin scaffolding ^101^. To suppress NMII activity, we used the blebbistatin family of class-II myosin ATPase inhibitors, conventional (-)-blebbistatin for endpoint experiments and (-)-4’-para-nitroblebbistatin for live imaging to avoid the phototoxicity associated with the parent compound (see Results for detailed pharmacological justification). To test the contribution of microtubule-dynein-based forces, we used dynarrestin, which binds the AAA3 site of the dynein motor domain, thereby inhibiting microtubule-stimulated ATPase and stepping activity ^106^. We favored dynarrestin over ciliobrevin-class dynein inhibitors, which additionally inhibit the proteasome and other AAA+ ATPases at effective concentrations, and over dynactin-disrupting strategies such as p150Glued CC1 dominant-negative expression or p50/dynamitin overexpression, which act at the level of cargo engagement rather than motor force generation.

Together, this pharmacological toolkit allowed us to functionally separate the contributions of three distinct cytoskeletal systems within the same live experiment, inhibiting dynein after oscillatory nucleokinesis was already established under UR214-9 and (-)-4’-para-nitroblebbistatin co-treatment, and directly testing whether dynein-dependent forces remained necessary once actomyosin contractility had been suppressed, a sequential perturbation design that would not have been feasible with a slower-onset genetic approach.

Transmigration through tight constrictions requires a “push-pull” cooperation between dynein and myosin II. Inhibiting either myosin II or dynein alone reduced transmigration by ∼40-45%, while combined inhibition nearly abolished it. This suggests that myosin II generates contractile force at the cell’s rear to build hydraulic pressure and push the nucleus ^17,100^, while dynein motors localized toward the cell front may pull the nucleus forward through an expanded microtubule network functioning as a force-transmission cable system. This cooperation is enabled by the reorganization of the microtubule network following septin inhibition, in which MTs expand, wrap around the nucleus, and extend into protrusions as active, load-bearing cables for long-range force transmission. Our computational model corroborates this framework, demonstrating that spontaneous polarization driving asymmetric dynein engagement, followed by nucleus-triggered repolarization, is sufficient to produce the sustained, self-organized nuclear oscillations observed experimentally. While our simplified model does not specify molecular interactions, mechanistically, nucleus-to-MT motor coupling likely involves the LINC complex of molecular adaptors, including SUN proteins at the inner nuclear membrane ^162,163^ and the KASH proteins at the outer nuclear membrane ^164–166^, anchoring the nucleus ^167^ to dynein motors ^168^ *via* Nesprin-2 ^5^ and BicD2 adaptors for the dynein complex ^5,169,170^. Nesprin1/2 may also couple with kinesin-1’s Kif5 *via* the LEWD motif ^171^ to assist nuclear movement ^172^, while dynein may interact with nuclear pore complexes *via* the RanBP2-BicD2 and Nup133-CENP-F adaptor complexes ^6,173^, facilitating nuclear movement, cell propulsion, and polarity maintenance ^174^.

Oscillatory nucleokinesis is not merely a structural feature of mesenchymally transformed T cells but constitutes the elementary mechanical unit of their migratory program. In UR214-9-treated hCD8⁺ T cells, sustained nuclear oscillations between competing protrusions resolve into net directional displacement through saltatory, stride-wise locomotion. Each stride is anchored by a discrete nuclear translocation event driven by dynein-generated forces transmitted along the microtubule network and anchored in protrusion-associated F-actin, completed by slingshot-like rear-protrusion detachment. This behavior is most pronounced in mature, exhausted hCD8⁺ T cells, consistent with their lower threshold for rear detachment and propensity for sustained mesenchymal spreading. The repeating mechanochemical cycle, consisting of protrusion extension, dynein engagement, nuclear translocation, and rear detachment, is fundamentally distinct from the pressure-driven, adhesion-independent peristalsis of amoeboid T cells or the characteristic single-stride nuclear translocation observed in developing neurons, and it extends the conceptual framework of dynein-driven nucleokinesis to adaptive immune cell motility.

Furthermore, pharmacological inhibition of dynein with Dynapyrazole A in zebrafish Langerhans cells produced a marked reduction in protrusion tip extension-retraction dynamics **(Figure 8)**, demonstrating that dynein-generated forces sustain cytoskeletal remodeling *in situ* within a physiologically native, mechanically confining epithelial environment. Because Langerhans cell protrusion dynamics are functionally analogous to the exploratory F-actin spreading observed in mesenchymally transformed T cells, this finding suggests that dynein-dependent force generation represents a shared cytoskeletal program across phenotypically distinct tissue-resident immune cell types, reflecting a broader, physiologically conserved mechanism for immune cell motility within the geometric constraints of solid tissues.

Collectively, these findings carry direct implications for cancer immunotherapy, where poor T cell trafficking into solid tumors remains a barrier to therapeutic efficacy. The dynein-microtubule force-transmission axis identified here operates in parallel with, and partially independently of, actomyosin contractility. The non-redundant cooperation between these systems during transmigration suggests that simultaneous targeting of both axes may yield greater therapeutic benefit than perturbing either system alone. While the current study relies heavily on septin inhibition as a tool to access the mesenchymal mechanotype, future studies should prioritize genetic dissection of the amoeboid-mesenchymal mechanotype continuum in physiologically relevant 3D environments to guide the engineering of CAR-T cells with self-sustained genetically enhanced deep-tissue penetration in solid tumor settings.

## METHODS

### Cell experiments

Primary human CD4^+^ and CD8^+^ T cells were isolated from whole human blood (STEMCELL Technologies Inc., USA, Cat# 70507.1) using EasySep Human CD4^+^ T Cell Isolation Kits (STEMCELL Technologies Inc., USA, Cat# 17952) and EasySep CD8⁺ T Cell Isolation Kit (STEMCELL Technologies Inc., catalog no 17953). Isolated CD4^+^ T cells were activated using ImmunoCult Human CD3/CD28/CD2 T cell activator and expanded in ImmunoCult-XF T cell expansion medium (STEMCELL Technologies Inc., USA, Cat# 10981) containing human recombinant interleukin-2 (IL-2; STEMCELL Technologies Inc., USA) at 37 °C in 5% CO2, per commercial protocol. CD8⁺ T cells were activated in complete RPMI-1640 medium (Gibco, 61870-036), supplemented with an additional 30 mL of human serum collected during isolation. Activation was carried out using ImmunoCult™ Human CD3/CD28 T Cell Activator (STEMCELL Technologies Inc., 10971) and IL-2. T Cell Activator was removed during expansion, and fresh IL-2 was provided to support optimal growth.

### Cell transmigration assay

Plasma-treated 3 µm Transwell inserts (Corning, Cat# 3415) were coated with 10 µg/mL recombinant human ICAM-1 protein (R&D Systems, Cat# 720-IC) for 1 hour at 37 °C and washed three times with PBS to remove excess ICAM-1. The wells of a 24-well plate were loaded with 1 mL of complete RPMI-1640 supplemented with 0.1 µg/mL human recombinant IL-2. The coated inserts were then placed into the wells, and 100 µL of complete RPMI-1640 medium was added to the insert, followed by the addition of activated CD8⁺ T cells (5*10^4^). After 6 hours of incubation at 37 °C in 5% CO_2_, the inserts were removed, and the transmigrated T cells in the lower chamber were quantified using a bright-field microscope.

### Prestained collagen gels

We premixed the gel on ice in a prechilled tube: 1 volume of 6 mg/mL collagen-1, e.g., CellAdhere type I (bovine, STEMCELL Technologies Inc., USA), 8/10 volume of ImmunoCult-XF T cell expansion medium, 1/10 volume of 10× phosphate-buffered saline (PBS; Thermo Fisher Scientific, Cat#10010023), 1/20 volume of 1 M Hepes, and 1/20 volume of 1 M Atto 647 N-hydroxysuccinimide ester (Sigma-Aldrich, Cat# 18373), or Alexa Fluor 568 ester (Molecular Probes, Cat# A20003), or Alexa Fluor 488 ester (Molecular Probes, Cat# A20000). A drop of gel premix (∼50 μL) was immediately spread onto the glass surface of a plasma-treated glass-bottom 35-mm petri dish (MatTek Corp., Cat# P35G-1.5-14-C) using a pipette tip. For polymerization, gels were covered with 1 mL of mineral oil (Sigma-Aldrich, Cat# M8410) to prevent water evaporation and incubated at room temperature overnight. Polymerized gels were rinsed with PBS to remove the unpolymerized gel components.

### Zebrafish husbandry

Zebrafish were housed at 26-27 °C on a 14/10 h light cycle. Animals aged 6-18 months of both sexes were used in this study. All zebrafish experiments were approved by the Institutional Animal Care and Use Committee at the University of Washington (Protocol #4439-01).

### Zebrafish scale removal

For scale removal, adult fish were anesthetized in system water containing 200 µg×mL^−1^ of buffered tricaine. Individual scales were removed with forceps, placed onto 6 mm plastic dishes, epidermis side up, and allowed to adhere for 30 s before adding L-15 medium (Gibco, USA, Cat# 21083027) prewarmed to room temperature (22 °C). Following scale removal, animals were recovered in system water.

### Dynapyrazole treatment, live-cell microscopy, and dendrite quantification of Langerhans cells

Zebrafish scales were pretreated for 2 hours with 11.5 µM dynapyrazole A (Sigma-Aldrich, USA, Cat# SML2127) or equivalent %v/v DMSO (Acros Organics, USA, Cat# 295522500) as vehicle control. Dishes were placed on the stage of an upright Nikon Ti-E microscope equipped with an A1R MP+ confocal scanner, a piezo *z*-drive (Mad City Labs Nano-F450), and a 25× water-dipping objective (1.1 NA; Nikon MRD77220). Confocal z-stacks were acquired every minute for 1 hour and post-processed using the denoise function in NIS-Elements.

To quantify dendrite dynamics, the ImageJ plugin MTrackJ ^175^ was used to manually track the dendrite tips of individual Langerhans cells at 1-minute intervals. The average speed of each dendrite was then averaged over the 1-hour period and plotted.

### Blebbistatin T cell treatment

To acutely inhibit non-muscle myosin-II-dependent actomyosin contractility during live imaging, T cells were treated with 50 µM (-)-4′-para-nitroblebbistatin (Cayman Chemical, USA, Cat# 24171), a photostable, low-phototoxicity blebbistatin derivative to acutely suppress NMII/class-II myosin motor activity under live-imaging conditions. Responses observed during live-cell treatment with (-)-4′-para-nitroblebbistatin were compared with parallel endpoint/control experiments performed using 50 µM (-)-blebbistatin (Sigma-Aldrich, USA, Cat# 203391). The 50 µM concentrations were chosen based on the reported rapid action in live cells ^114^ and biochemical measurements showing suppression of myosin-II ATPase activity ^113,115^.

### Dynarrestin T cell treatment

Cytoplasmic dynein (dynein-1) activity was acutely inhibited with dynarrestin (AAA3-site motor domain inhibitor; Tocris-Biotechne, USA, Cat# 6525 ^106^). Dynarrestin solution was added directly to the imaging medium to a final working concentration of 50 µM during live time-lapse acquisition. Dynarrestin was used in place of genetic depletion to avoid confounding effects of cytoskeletal remodeling associated with chronic loss of dynein activity and to permit within-cell before/after comparisons, given its acute (minutes-scale) onset.

### UR214-9 T cell treatment

Septin GTPase activity was acutely inhibited using UR214-9, a chlorinated forchlorfenuron (FCF) analog developed through structure-activity relationship optimization of the parent compound and selected for substantially greater potency than FCF ^104^. UR214-9 was added directly to the imaging media at a final concentration of 50 µM during live time-lapse acquisition to induce amoeboid-to-mesenchymal mechanotype transition ^101^ and oscillatory nucleokinesis. UR214-9 was sourced from Dr. Rakesh Singh (University of Rochester Medical Center, USA) and then from Focus Biomolecules (USA, Cat# 10-4397).

### Rho Activator II T cell treatment

RhoA, -B, and -C were activated with Rho Activator II (Cytoskeleton Inc, USA, Cat# CN03), a derivative of bacterial CNF toxin, which deamidates glutamine-63 in the Switch II region and prevents GTP hydrolysis. Rho Activator was directly added to the medium at a manufacturer-recommended concentration of 1.0 µg/ml during the 6-hour transmigration experiment ^176^.

### Rho Inhibitor I T cell treatment

RhoA, -B, and -C were inhibited with Rho Inhibitor I (Cytoskeleton Inc, USA, Cat# CT04), a derivative of Clostridium botulinum C3 Transferase, which ADP-ribosylates asparagine 41 in the GTPase effector-binding domain and prevents Rho signaling. Rho Inhibitor was added directly to the medium at a recommended concentration of 1.0 µg/ml during the 6-hour transmigration experiment ^177^.

### Calyculin A T cell treatment

To induce actomyosin contractility during the transmigration, T cells were treated with 1.0 ng/ml a marine-derived toxin, Calyculin A (Sigma Aldrich, USA, Cat. #C5552), a potent protein myosin light chain phosphatase (MLCP) inhibitor, which enhances myosin II phosphorylation and actomyosin contractility ^178^.

### T cell microscopy and image processing

Tissue and cell samples were fixed with 4% paraformaldehyde (PFA; Sigma-Aldrich, Cat# P6148) in PBS overnight at 4 °C or for 30 min at room temperature. PFA-fixed tissues were rinsed with PBS, placed in 30% sucrose, and then cryosectioned using a Leica CM1860. The 100-micron-thick sections were stored in PBS containing 0.1% sodium azide (Sigma-Aldrich, Cat# 2002) at 4 °C until further processing. For immunostaining, PFA-fixed samples were rinsed with 1% bovine serum albumin (BSA; Thermo Fisher Scientific, Cat# BP9704) in PBS, permeabilized with 0.1% Triton X-100 (Sigma-Aldrich, Cat# X100) in PBS for 30 min, and blocked with 1% BSA in PBS for 60 min. For staining, all primary antibodies were diluted in 1% BSA in PBS. The incubation duration with any of the listed primary antibody solutions was 2 hours at room temperature. Primary antibodies were: monoclonal Rat IgG2b Alexa Fluor® 647 anti-mouse CD3 Antibody (BioLegend, Cat# 100209), Rat alpha-Tubulin Monoclonal Antibody YL1/2 (Thermo Fisher Scientific, Cat# MA1-80017). Catalog numbers for these reagents and other key resources are listed in the Key Resources Table. Labeling with Alexa Fluor-conjugated secondary antibodies (Thermo Fisher Scientific) was performed at a final concentration of 5 μg/mL for 1 hour in 1% BSA in PBS at room temperature. After washing out excess secondary antibodies, chromatin and actin, if necessary, were labeled with 1:1000 Hoechst solution (Tocris, Cat#5117) and SiR-actin (Cytoskeleton Inc., Cat# CY-SC001) or phalloidin conjugates (Sigma-Aldrich, Cat# 49409, Thermo Fisher Scientific, Cat# A12380), respectively. Samples were mounted for imaging in 90% glycerol (Sigma-Aldrich, Cat# G5516) in 1×PBS.

Imaging was performed using an instant structured illumination microscopy (iSIM) system (VisiTech Intl, Sunderland, UK) equipped with an Olympus UPLAPO-HR ×100/1.5 NA objective, Zeiss LSM 980 Airyscan 2 (Zeiss Microscopy, Oberkochen, Germany) with Plan-Apochromat 63×/1.4 Oil objective, and Leica Microsystems TCS SP8 laser-scanning confocal microscope using LIAchroic Lightning system and LAS-X Lightning Expert deconvolution software. The default optical configuration used for 3D imaging was: a 40×/1.3 NA oil-immersion objective (Leica, Germany), HyD 2.0-SMD detectors, a 0.85 AU pinhole, and a scanning frequency of 100 Hz. Acquisition settings were optimized *via* Nyquist LAS-X. Fluorescence channels were acquired sequentially to prevent excitation and emission signal bleeding between channels. Live-cell imaging was performed on a widefield epifluorescence Leica DMi8 or Leica SP8 microscope system (Leica, Germany) housed within an environmental chamber maintaining 37 °C, 5% CO_2_, and controlled humidity. Brightfield and fluorescence time-lapse images were acquired at 20×, 40×, and 63× magnifications using LAS-X software (Leica, Germany).

Images were deconvolved with an iSIM-specific commercial plugin from Microvolution (Cupertino, CA) in FIJI and Zeiss Zen Blue software (Oberkochen, Germany). Analyses were performed using Leica LAS-X software (Leica, Germany) and ImageJ/Fiji. Figures were composed using unmodified LAS X- and Zeiss Zen Blue-generated images with Adobe Illustrator CC 2024 (Adobe).°C

### High-precision micropatterning

ICAM-1 micropatterning was performed using indirect, high-definition, submicron-scale polyacrylamide gel (PAA) micropatterning, adapted from a previously established protocol ^179^. To prevent microstamp collapse, we used a composite microstamp consisting of a 0.5-0.8 mm-thick hard PDMS (hPDMS) veneering microprinting surface mounted on a 5-8 mm-thick cushioning layer of regular PDMS (rPDMS) blocks ^37,180^.

The hPDMS mixture was prepared by thoroughly mixing 3.4 g of VDT-731 (Gelest, Inc., Cat# VDT-731), 18 µL of Pt catalyst (Sigma-Aldrich, Cat# 479543), and one drop of cross-linking modulator (Sigma-Aldrich, Cat# 396281). Before use, 1 g of HMS-301 (Gelest, Inc., Cat# HMS-301) was added, mixed by vortexing for 30 seconds, and then degassed in a high-speed centrifuge (3 minutes at 3000 rpm, 4 °C). To prevent hPDMS from curing before use, the mix must be kept cold on ice.

For microstamp preparation, molds (Minnesota Nano Center, University of Minnesota) were coated with a 0.5-0.8 mm layer of hPDMS and cured at 70 °C for 30 minutes. An ∼8 mm-thick layer of bubble-free rPDMS (Sylgard-184, 1:5 curing agent/base ratio) was cured atop the hPDMS layer (70 °C for ∼1 hour). The cured composite was then peeled and cut into 1 × 1 cm pieces.

PAA premixes used a 40% acrylamide (40% AA) base (BioRad, Cat# 161-0140) and a 2% bis-AA cross-linker (BioRad, Cat# 161-0142) ^181,182^. Streptavidin-acrylamide (Thermo Fisher, Cat# S21379) was added to a final concentration of 0.133 mg/mL to cross-link PAA gels with biotinylated proteins. For 50 µL of G’ = 50 kPa PAA gel premix, components were mixed as follows: 40% AA: 15 µL, 2% bis-AA: 14.40 µL, 2 mg/mL streptavidin-acrylamide: 3.33 µL, 10× PBS: 5 µL, deionized Milli-Q water: 11.17 µL, TEMED: 0.1 µL.

The premix solutions were degassed and stored at 4 °C. Curing was initiated by adding 1 µL of 10% APS to 50 µL of PAA premix immediately before casting.

ICAM-1 pattern assembly order:

1. **Intermediate Printing:** The microstamps were coated with 0.2 mg/mL anti-Human-Fc Fab antibody fragments (Jackson Immunoresearch, Cat# 109-007-008, dissolved in PBS) for 40 min at 37 °C. The stamps were then rinsed in deionized water and thoroughly dried under argon or nitrogen jet immediately prior to soft lithography. For crosslinking to PAA and fluorescence, the FAB against ICAM-1-hFc was covalently conjugated to biotin ((+)-biotin *N*-hydroxysuccinimide ester, Sigma-Aldrich, Cat# H1759) and a fluorescent AF488 NHS ester (Lumiprobe, Cat#11820), following the manufacturer’s protocols.
2. **Glass Activation:** To ensure covalent cross-linking of PAA to glass-bottomed inserts in 35 mm cell culture dishes (MatTek Corp., Cat# P35G-1.0-20-C), the glass surfaces were activated with 3-(trimethoxysilyl)propyl methacrylate (Sigma-Aldrich, Cat# 6514) according to the manufacturer’s protocols.
3. **PAA Casting:** 5 µL of PAA premix (G’ = 50 kPa, see **PAA Elastic Gel Premix**) containing 5% streptavidin-acrylamide (ThermoFisher, Cat# S21379) was ‘sandwiched’ between the activated glass surface and the micropatterned intermediate glass surface immediately after adding the curing catalyst (ammonium persulfate, APS).
4. **Pattern Transfer:** The PAA sandwiches were incubated in deionized water (1 h at 20 °C) to induce hypotonic swelling and gentle coverglass release.
5. **Protein Incubation:** The resulting PAA gels were incubated with 10 µg/mL ICAM1-hFc solution (R&D Systems, Cat# 720-IC) in cold PBS (4 °C, 12 hours), rinsed three times with cold sterile PBS, and used in experiments.

### Flow cytometry analysis of microtubule density in live cells

For analysis of microtubule density in live cells, we live-stained control (DMSO)-treated and UR214-9-treated (25 µM) CD4^+^ and CD8^+^ T cells with SiR-Tubulin, following the commercial protocol, at 37 °C in 5% CO2. Briefly, a cell aliquot was separated into eight tubes as follows: unstained control and UR214-9-treated cells, control and UR214-9-treated cells stained with propidium iodide (PI, 0.5mg/mL, 1:200, for 10 minutes prior to flow cytometry), control and UR214-9-treated cells stained with the SiR-reagent, and control and UR214-9 cells stained both with PI and SiR-reagent. All UR214-9-treated cells were incubated with UR214-9 for 1 hour before the addition of SiR-reagent, and for an additional hour after the addition of SiR-reagent. After the second hour of incubation, we added PI as necessary and proceeded with flow cytometry analysis.

### Computational Model of Cell Morphodynamics and Nuclear Dynamics

We developed an integrative mechanochemical model of cell morphodynamics that couples protrusive cell edge dynamics with intracellular signaling and force generation. The model builds upon our previously described framework ^136^ and extends it by explicitly incorporating the nucleus and dynein-mediated force transmission.

### Cell Representation and Edge Dynamics

The cell is represented as a binary domain *m_i_*_,*j*_(*t*) ∈ (0, 1) on a square lattice, where shape evolution is governed by stochastic protrusion and retraction events at the cell outline. At each iteration, candidate pixels at the edge are selected for protrusion or removal based on the following probabilities:

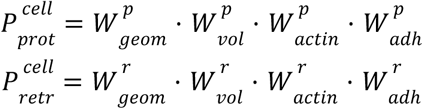

where:

- 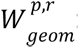: local curvature-dependent geometry factor
- 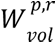: cell volume regulation factor
- 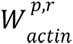: actin regulation factor
- 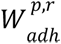: adhesion presence factor

The geometry factor depends on local curvature computed from the neighborhood of each boundary pixel (*i*, *j*), such that convex regions favor retraction and concave regions favor protrusion:

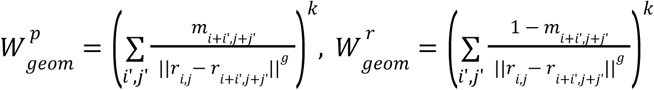

where *r_i_*_,*j*_ is the pixel position, (*i*’, *j*’) are indices of the neighboring pixels, *g* is the sensitivity to curvature, and *k* controls the contribution of this factor to the total probability.

The volume factor maintains overall cell size:

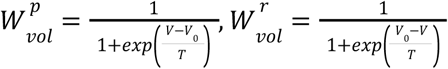

where *V* is the current 2D cell area, *V*_0_ is the target (or initial) area, and *T* is the sensitivity to the deviation of the area from its target value.

The actin factor depends on local nucleation-promoting factor (NPF, e.g., Arp2/3) or GTPase (e.g., Rac1 or Cdc42) activity:

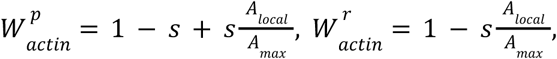

where *A_local_* is the mean concentration over the pixel neighborhood of the active form of an NPF or GTPase, *A_max_* is the maximal concentration in the cell, and *s* is the slope of the probability dependence on the activity level.

The adhesion factor is defined by the adhesion field, which is calculated from the mask of adhesion locations *r_n_*, smoothed by a Gaussian filter, and scaled to have values between 0 and 1:

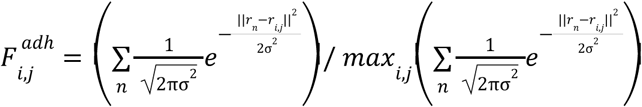

where *r_n_* is the position of adhesion *n* and σ is the standard deviation of the Gaussian function. The resulting adhesion factor becomes:

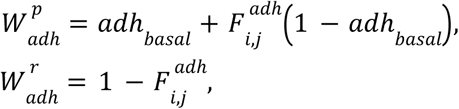

where *ad*ℎ*_basal_* is the basal (minimal) contribution to protruding in the absence of nearby adhesions. This formulation ensures that the cell cannot retract past an adhesion point.

Cell shape updates are performed using a Monte Carlo scheme that alternates protrusion and retraction steps (every *diff_T_* time step of the reaction-diffusion solver), thereby controlling the protrusion-retraction time scale relative to the activation time scale of actin regulation. The algorithm prohibits protrusion-retraction events that violate cell connectivity or lead to hole formation. Throughout the simulation (after every *ad*ℎ*_T_* time step), a specified fraction *ad*ℎ*_frac_* of the total number of adhesions *ad*ℎ*_tot_* is randomly rearranged within the moving cell, and the adhesion field is recalculated accordingly.

### Explicit Nuclear Representation and Dynein Forces

To capture nucleokinesis, we introduced two extensions:

#### Nucleus

The nucleus is represented as an internal deformable domain with its own geometry and rigidity constraints. Its shape evolves via the same protrusion–retraction framework, with additional constraints favoring compact geometry (global geometry factor) and the influence of dynein forces:

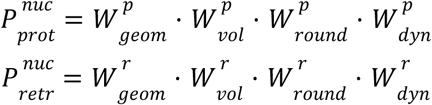

with 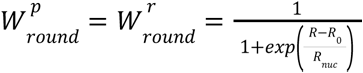, where *R* is the current roundness of the 4-connected nucleus shape (perimeter^2^/area), *R*_0_ is the target roundness, and *R_nuc_* is the sensitivity to deviation from a compact (round) shape.

#### Dynein-Mediated Pulling Field

Dynein forces are incorporated as an effective pulling field acting on the nucleus. This field is constructed by mapping protrusion geometry to force direction:

1. For each cell edge pixel, the distance to the nearest nuclear boundary point (*i*. *j*) is computed.
2. The concentration of the component engaging dynein at each cell edge pixel is mapped to the nearest nuclear boundary point (minimal distance) and accumulated into the field 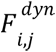.
3. Thus, regions of the nuclear periphery aligned with a higher concentration of the polarization component (see the RD model) produce larger values of 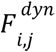 and, consequently, stronger pulling forces.
4. In a variation of the model assuming a uniform distribution of engaged dyneins along the protrusion length, the field 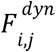 is calculated from contributions proportional to the product of the polarization component concentration and the distance between the cell and nuclear edges. Thus, in this model, the pulling force depends on both the dynein engagement level and the length of the protrusions where that engagement occurs.
5. After accounting for the contribution from all points of the cell edge, the field 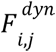 is normalized to the (0,1) range.

The dynein contribution modifies nuclear protrusion/retraction probabilities as:

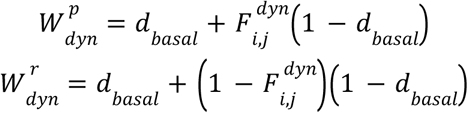

where *d_basal_* is the basal (minimal) contribution of dynein factor to the total probability of the pixel-size protrusion-retraction events defining nuclear shape changes and its translocation along cell protrusions.

### Reaction–Diffusion Model of Regulatory Components

Intracellular signaling is modeled using a mass-conserved reaction–diffusion (MCRD) system representing active (*A*) and inactive (*I*) NPF or GTPase, coupled to F-actin (*F*):

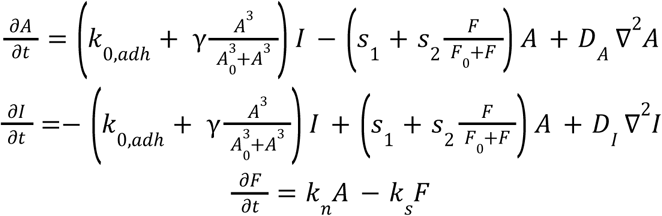

This formulation closely follows the functional and parameter choices of the model originally studied by Holmes *et al.* ^183^, except that it is here coupled to cell shape (i.e., solved on a moving 2D domain) and to the adhesion distribution that influences the activation rate. The model includes

- *A*: active NPF or GTPase (membrane-bound)
- *I*: inactive NPF or GTPase (cytosolic)
- *F*: F-actin
- *D_A_*, *D_I_*: diffusion coefficients (*D_A_*≪ *D_I_*)
- ∇^2^: the 2D Laplacian operator
- autocatalysis: Hill term
- negative feedback: F-actin-mediated suppression
- No-flux (Neumann) boundary condition.

The activation rate of the NPF is modulated spatially by adhesion locations as:

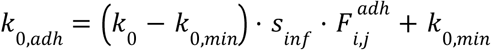

where *k*_0,*min*_ is the minimal value of the basal activation rate in the absence of the adhesion influence, *k*_0_ is the basal activation rate at immediate proximity to adhesions, and *s_inf_* modulates the strength of the adhesion effect. For GTPase dynamics responsible for polarity establishment, we assumed no direct adhesion modulation *s_inf_* = 0 and *k*_0,*min*_ = *k*_0_. This system produces excitable and oscillatory regimes consistent with observed polarity dynamics.

In a variation of the model assuming delayed dynein engagement, we included a relatively slow component, responsible for dynein engagement and positively regulated by the active GTPase:

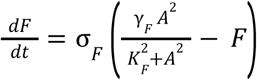

### Coupling Between Mechanics and Signaling

The bidirectional coupling of the model components produces the following relationships:

- NPF or GTPase activity and adhesion distributions (RD + AB modules) → F-actin polymerization → protrusion/retraction modulation → cell shape change
- Cell shape → local curvature → polarity localization
- Actin polarization → dynein engagement → nuclear motion
- Nuclear position → local cytoplasmic exclusion → RD reset

This feedback generates cyclic polarity switching and oscillatory nucleokinesis.

### *Numerical* Implementation

- Spatial discretization: square lattice (pixel-based)
- Time integration: forward Euler for the RD system
- Alternating update scheme:

- protrusion step
- retraction step
- RD update
- Stochastic adhesion redistribution

Time step and grid spacing satisfy the diffusion stability criterion:

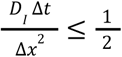

Mass conservation and positivity constraints are enforced at each step.

## Parameter Table

| Parameter | Description | Value |
| --- | --- | --- |
| $k$ | Geometry factor weight for the cell edge | 1.6 |
| $k_{nuc}$ | Geometry factor weight for the nucleus edge | 4 |
| $g$ | Curvature sensitivity | 2 |
| $T$ | Volume sensitivity for the cell | 500 |
| $T_{nuc}$ | Volume sensitivity for the nucleus | 1 |
| $s$ | Actin factor slope | 0.03 |
| $\sigma$ | STD of Gaussian smoothing of the adhesion field | 5 |
| $adh_{basal}$ | Basal contribution of adhesion field to protrusion | 0.3 |
| $adh_{tot}$ | Total number of adhesions | 50 |
| $adh_{frac}$ | Fraction of adhesions that are rearranged at each $adh_T$ timestep | 0.03 |
| $R_0$ | Targeted nuclear roundness (4-connected shape) | 20 |
| $R_{nuc}$ | Sensitivity to changes in roundness | 1 |
| $d_{basal}$ | Basal contribution of dynein force field to nuclear motion | 0.9 |
| $D_A$ | Diffusion coefficient (active NPF or GTPase) | 0.001/3 |
| $D_I$ | Diffusion coefficient (inactive NPF or GTPase) | 0.1/3 |
| $k_0$ | Basal activation rate near adhesions | 0.1 |
| $k_{0,min}$ | Minimum activation rate (far from adhesions) | 0.01 |
| $s_{inf}$ | Strength of the influence of adhesions on activation | 10 |
| $\gamma$ | Autocatalytic strength | 1 |
| $A_0$ | Hill threshold in the autocatalytic feedback | 0.4 |
| $s_1$ | Basal deactivation rate | 0.7 |
| $s_2$ | Negative feedback strength | 0.20 (for NPF)<br>0.06 (for GTPase) |
| $F_0$ | Sensitivity of negative feedback to F-actin | 0.5 |
| $k_n$ | F-actin polymerization rate | 0.1 |
| $k_s$ | F-actin degradation rate | 0.025 |
| $K_F$ | Sensitivity of the slow component to the GTPase | 0.4 |
| $\sigma_F$ | Turnover rate scale for the slow component | 0.001 |
| $\gamma_F$ | Strength of GTPase influence on the slow component | 1 |
| $\Delta t$ | Temporal step of simulation (RD solver) | 0.000375 |
| $\Delta x$ | Spatial step of simulation (pixel size) | 0.0070755 |
| $diff_T$ | Number of RD iterations between shape (protrusion/retraction) updates | 100 |
| $adh_T$ | Number of RD iterations between adhesion updates | 200 |

## Key resources table

| REAGENT or RESOURCE | SOURCE | IDENTIFIER |
| --- | --- | --- |
| **Antibodies** |  |  |
| AffiniPure Fab Fragment Goat Anti-Human IgG, Fc<sub>γ</sub> fragment specific | Jackson ImmunoResearch Laboratories, Inc | Cat# 109-007-008; RRID:AB_2632440 |
| Anti-Tubulin antibody [YL1/2] | AbCam | Cat# ab6161; RRID:AB_305329 |
| Anti-Tubulin antibody [SAP.4G5] | Sigma-Aldrich | Cat# T7816, RRID:AB_261770 |
| Anti-Tubulin Chicken Polyclonal Antibody | VWR | Cat# 89416-948 |
| Recombinant Anti-alpha Tubulin antibody [EP1332Y] - Microtubule Marker | AbCam | Cat# ab52866 |
| Atto 647N goat anti-mouse IgG | Sigma-Aldrich | Cat# 50185, RRID:AB_1137661 |
| Goat anti-Mouse IgG, Cross-Adsorbed Secondary Antibody, Alexa Fluor™ 555 | Invitrogen | Cat# A-21127 |
| Goat anti-Chicken IgY (H+L) Secondary Antibody, Alexa Fluor™ 488 | Invitrogen | Cat# A-11039 |
| Anti-Rabbit-IgG - Atto 647N antibody produced in goat | Sigma-Aldrich | Cat# 40839 |
| Alexa Fluor 488 donkey anti-rat IgG (H+L) | Thermo Fisher | Cat# a21208; RRID:AB_141709 |
| **Chemicals, Peptides, Recombinant Proteins, and Consumables** |  |  |
| HMS-31, (25-35% Methylhydrosiloxane)-Dimethylsiloxane Copolymer, Trimethylsiloxane Terminated | Gelest, Inc | Cat# HMS-301; CAS#68037-59-2 |
| VDT-731, (7.0-8.0% Vinylmethylsiloxane) - Dimethylsiloxane Copolymer, Trimethylsiloxyl Terminated | Gelest, Inc | Cat# VDT-731; CAS#67762-94-1 |
| 2,4,6,8-Tetramethyl-2,4,6,8-tetravinylcyclotetrasiloxane | Sigma-Aldrich | Cat# 396281; CAS#2554-06-5 |
| Platinum(0)-2,4,6,8-tetramethyl-2,4,6,8-tetravinylcyclotetrasiloxane complex solution | Sigma-Aldrich | Cat# 479543; CAS#68585-32-0 |
| SYLGARD 184 Silicone Elastomer Kit, 0.5 kg KIT | Dow Corning, Sigma-Aldrich | Cat# 4019862; CAS#68988-89-6 |
| 40% Acrylamide Solution, Electrophoresis purity reagent, 500 mL | BioRad | Cat# 161-0140 |
| 2% Bis Solution, 500 mL | BioRad | Cat# 161-0142 |
| Streptavidin Acrylamide, 1 mg | Thermo Fisher, Life Technologies | Cat# S21379 |
| TEMED | Thermo Scientific | Cat# 17919; CAS#110-18-9 |
| Ammonium Persulfate, BioUltra, for molecular biology | Fluka Analytical | Cat# 09913-100G; CAS#7727-54-0 |
| 3-(Trimethoxysilyl)propyl methacrylate | Sigma-Aldrich | Cat# 6514; CAS#2530-85-0 |
| Ethyl Alcohol 200 Proof, Absolute, Anhydrous ACS/USP Grade | Pharmco-Aaper | Cat# 111000200; CAS#64-17-5 |
| Collagen-1, rat | Corning, NY | Cat# 354249 |
| Collagen-1, bovine | Sigma-Aldrich | Cat# 804622 |
| Genemate LE Quick Dissolve Agarose | BioExpress | Cat# E-3120-500FG |
| ICAM1, Human Protein, Recombinant, hlgG1-Fc.His Tag, Active | Sino Biological, China | Cat# 10346-H03H-50 |
| Paraformaldehyde, reagent grade, crystalline | Sigma-Aldrich | Cat# P6148-500G; CAS#30525-89-4 |
| Triton X-100 | Sigma-Aldrich (Roche) | Cat# 11332481001 CAS#9002-93-1 |
| DMSO (Dimethyl sulfoxide) | Sigma-Aldrich | Cat# 472301-100ML; CAS#67-68-5 |
| (-)-4' *para*-nitroblebbistatin. | Cayman Chemical | Cat# 24171; CAS#1621326-32-6 |
| (-)-Blebbistatin | Sigma-Aldrich | Cat#203391 |
| Dynarrestin | Tocris-Biotechnne | Cat# 6525 |
| Dynapyrazole A | Sigma-Aldrich | Cat# SML2127 |
| UR214-9 | Dr. Rakesh Singh | Custom synthesis |
| UR214-9 | Focus Biomolecules | Cat# 10-4397 |
| 0.25% Trypsin, 2.21 mM EDTA, 1× [-] sodium bicarbonate | Corning | Cat# 25-053-CI |
| Plasmocin | Fisher Scientific | Cat# NC9698402 |
| Penicillin Streptomycin Solution, 100X | Corning | Cat# 30-002-CI |
| Fetal Bovine Serum | HyClone | Cat# SH30910.03 |
| Acetic Acid, Glacial | Fisher Chemical | Cat# BP2401-500; CAS#64-19-7 |
| 35 mm Dish \| No. 1.0 Coverslip \| 20 mm Glass Diameter \| Uncoated | MatTek Corporation | Cat# P35G-1.0-20-C |
| WillCo-dish Glass Bottom Dish \| No. 1.5H Coverglass thickness \| 40 mm Glass Diameter \| Uncoated | WillCo Wells B.V. | Cat# HBST-5040 |
| Cover Glasses, FisherFinest Rectangles, 25×60-1, Thickness 0.13-0.17 mm | Fisher Scientific | Cat# 24X60-1 |
| Cover Glasses, Circles, 15 mm, Thickness 0.13-0.17 mm | Carolina Biological Supply Company | Cat# 633031 |
| SiR Tubulin, Spirochrome Kit | Cytoskeleton Inc. | Cat# CY-SC002 |
| Phalloidin-Atto 488 | Sigma-Aldrich | Cat# 49409 |
| Phalloidin-iFluor 647 Reagent - CytoPainter | AbCam | Cat# ab176759 |
| Bovine Serum Albumin (BSA), fatty acid-free powder | Fisher Bioreagents | Cat# BP9704-100; CAS#9048-46-8 |
| Propidium iodide, 0.5 mg/ mL | Sigma-Aldrich | Cat# 11 348 639 001 |
| (+)-Biotin N-hydroxysuccinimide ester | Sigma-Aldrich | Cat# H1759; CAS#35013-72-0 |
| Alexa Fluor 568 succinimidyl ester (NHS) | Molecular Probes | Cat# A20003 |
| Alexa Fluor 488 succinimidyl ester (NHS) | Molecular Probes | Cat# A20000 |
| Alexa Fluor 405 succinimidyl ester (NHS) | Molecular Probes | Cat# A30000 |
| Atto 647 NHS ester | Sigma-Aldrich | Cat# 07376 |
| Hoechst 33342, Fluorescent Dye for labeling DNA | Tocris | Cat# 5117; CAS#23491-52-3 |
| DRAQ5 Fluorescent Probe Solution (5 nM) | Thermo Fisher Scientific | CAT# 62251 |
| Slide-A-Lyzer MINI Dialysis Device, 7K MWCO, 0.1 mL | Thermo Fisher | Cat# 69560 |
| 6.5 mm transmigration TransWell inserts with 3 μm-pores in 10-μm-thick polycarbonate membranes | Costar | Cat# 3414 |
| Rho Activator II | Cytoskeleton, Inc. | Cat# CN03 |
| Rho Inhibitor I | Cytoskeleton, Inc. | Cat. #CT04 |
| Calyculin A | Sigma-Aldrich | Cat. #C5552 |
| Hepes (1M) | Thermo Fisher Scientific | Cat# 15630080 |
| Phosphate Buffered Saline (PBS) pH 7.4 (1X) | Thermo Fisher Scientific | Cat# 10010023 |
| Penicillin Streptomycin Solution, 100X | Corning | Cat# 30-002-CI |
| RPMI 1640 with L-glutamine | Corning | Cat# 10-040-CV |
| EasySep Human CD4+ T Cell Isolation Kit | STEMCELL Technologies, Inc., USA | Cat# 17952 |
| EasySep Human CD8+ T Cell Isolation Kit | STEMCELL Technologies, Inc., USA | Cat# 17953 |
| ImmunoCult Human CD3/CD28 T Cell Activator | STEMCELL Technologies Inc., USA | Cat# 10971 |
| "The Big Easy" EasySep Magnet | STEMCELL Technologies, Inc., USA | Cat# 18001 |
| Lymphoprep | STEMCELL Technologies, Inc., USA | Cat# 18060 |
| L-glutamine (200mM) | Gibco | Cat# 25-030-081 |
| EmbryoMax Penicillin-Streptomycin Solution, 100X | Sigma-Aldrich | Cat# TMS-AB2-C |
| ImmunoCult-XF T Cell Expansion Medium | STEMCELL Technologies, Inc., USA | Cat# 10981 |
| Human Recombinant IL-2, ACF | STEMCELL Technologies, Inc., USA | Cat# 78145.1 |
| DMSO | Acros Organics | 295522500 |
| L-15 Media | Gibco | 21083027 |
| **Experimental Models: Cell Lines** |  |  |
| Human CD4+ T cells<br>(derived from commercially supplied human whole peripheral blood). | STEMCELL Technologies, Inc.,<br>USA | Cat#70507.1 |
| Human CD8+ T cells (derived from commercially supplied human whole peripheral blood). | STEMCELL Technologies, Inc.,<br>USA | Cat#70507.1 |
| **Experimental Models: Zebrafish** |  |  |
| *Tg(mpeg1.1:YFP)<sup>w200tg</sup>* | Roca and Ramakrishnan, 2013 <sup>184</sup> | ZFIN: ZDB-TGCONSTRUCT 130130-3 |
| *Gt(ctnna-citrine)<sup>Ct3a</sup>* | Zigman et al., 2011 <sup>185</sup> | ZFIN: ZDB-ALT-111010-23 |
| **Software and Algorithms** |  |  |
| KaleidaGraph 5 for MacOS | http://www.synergy.com/ | RRID:SCR_014980 |
| GraphPad Prism 7b | https://www.graphpad.com/ | RRID:SCR_002798 |
| PlotsOfData | https://huygens.science.uva.nl | N/A |
| Adobe Illustrator CC, 21.0.0. | Adobe Systems, Inc. | RRID:SCR_010279 |
| MATLAB R2019a | The MathWorks |  |
| LAS X | Leica Microsystems |  |
| Fiji | https://imagej.net/software/fiji/ |  |
| ZEN | Zeiss Microscopes |  |
| **Other** |  |  |
| Leica SP8 laser scanning confocal microscopy imaging system | Leica Microsystems<br>https://www.leica-microsystems.com/ |  |
| TokaiHit Stage Top Incubator, Model STXG-GSI2X-SET | Tokai Hit, Fujinomiya, Japan |  |
| Zeiss LSM 980 Airyscan 2 imaging system | Zeiss Microscopy, Oberkochen, Germany |  |
| Instant structured illumination (iSIM) microscopy imaging system | VisiTech Intl, Sunderland, UK |  |
| Nikon Ti-E microscope with A1RMP+ confocal scanner | Nikon, Japan |  |

## Data Availability

All data supporting the findings of this study are available within the manuscript. All other data supporting the findings of this study can be obtained from the corresponding authors upon reasonable request. Source data will be available through the Dryad Digital Repository.

## Code Availability

The source code, configuration files, and analysis scripts used to generate the computational results presented in this study will be publicly available through the Dryad Digital Repository.

## Acknowledgements

This research was supported in part by the Intramural Research Program of the National Institutes of Health (NIH), National Institute of Biomedical Imaging and Bioengineering Division of Intramural Research with grant number ZIA-EB000094 to A.X.C.R and Y.T, LEO Foundation award LF-OC-24-001646 to J.P.R., and by U.S. Army Research Office (ARO), award No. W911NF261A112 to D.T. The contributions of the NIH authors were made as part of their official duties as NIH federal employees, are in compliance with agency policy requirements, and are considered Works of the United States Government. A.Z. efforts were supported by the FDA Intramural Research Program of the Center for Biologics Evaluation and Research. The findings and conclusions presented in this paper are those of the authors and do not necessarily reflect the views of the NIH, FDA, or the US Department of Health and Human Services. E.T. efforts in this research project are supported byThe Penn State University startup funds.

## AUTHORS CONTRIBUTIONS

Y.T., D.T., A.S.Z., and E.D.T. conceptualized the work and developed the theoretical and experimental approaches. D.T. designed and executed the nucleokinesis model. E.P. and J.R. provided quantified experimental data on zebrafish. Y.T., N.S., A.X.C.R., A.S.Z., and E.D.T. performed human T cell experiments and analyzed the data. Y.T., N.S., D.T., A.S.Z., and E.D.T. wrote the manuscript, with editorial contributions from all authors. D.T. and E.D.T. oversaw all aspects of the study.

## SUPPLEMENTARY INFORMATION

**Supplemental Figure 1.**
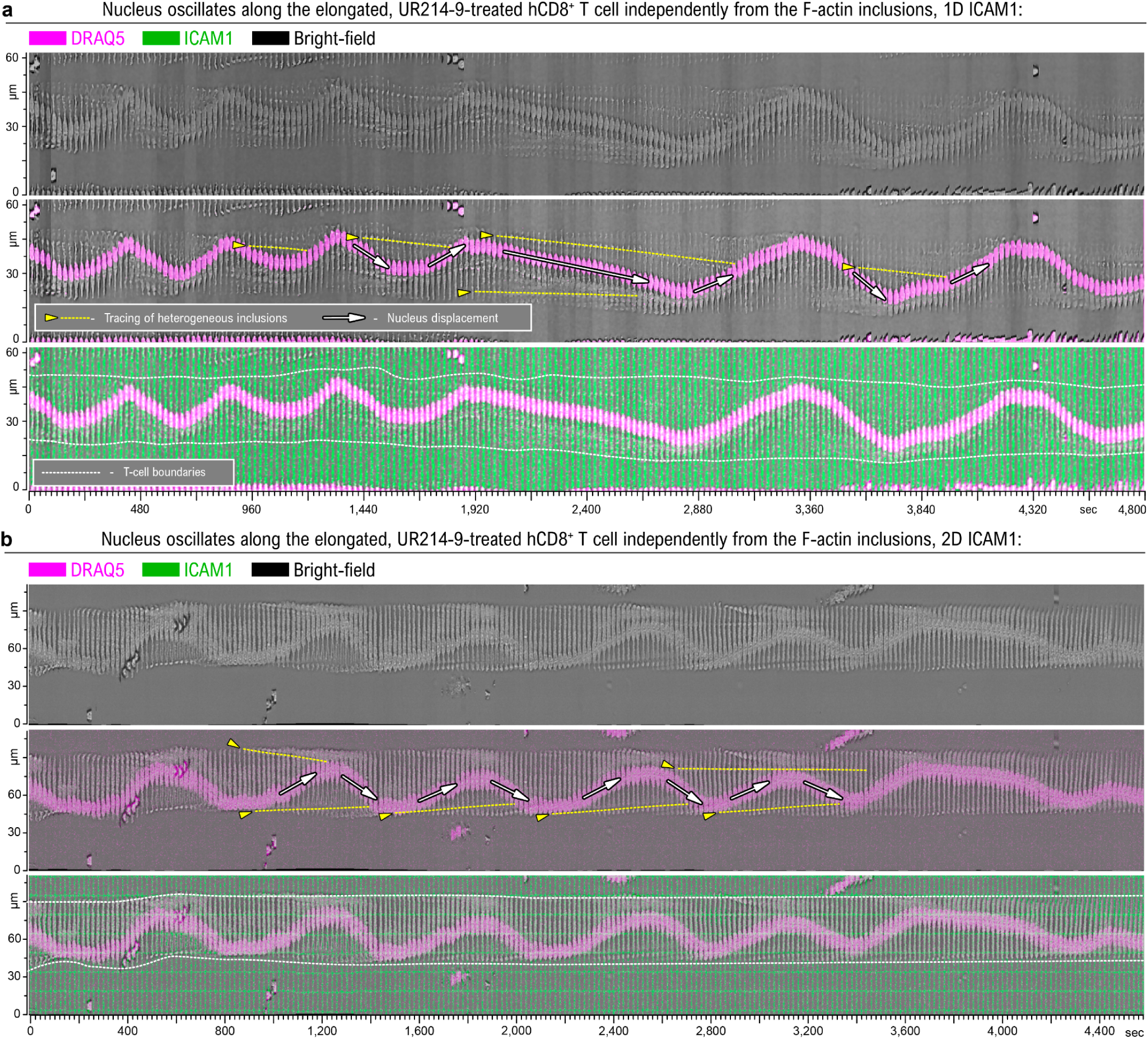
Nuclear oscillation is decoupled from F-actin heterogeneity dynamics in elongated UR214-9-treated hCD8^+^ T cells. **a** - Kymograph of nuclear oscillations in a human primary CD8^+^ T cell along the 1D ICAM1 1 µm-wide lane. Fluorescence imaging of the oscillating nucleus (*magenta*, DRAQ5-labeled) and the underlying adhesive 1 µm-wide ICAM1 lane (*green*). *Note the linear (centripetal) displacement of F-actin heterogeneities within the elongated T cell body (dashed lines), as the nucleus completes a full oscillatory back-and-forth cycle.* See **Movie 9.** **b** - Kymograph of nuclear oscillations in a human primary CD8^+^ T cell spread along the 2D ICAM1 grid (1 µm-wide lanes, 15 µm-pitch, crisscrossed orthogonal grid). *Note the linear (centripetal) displacement of F-actin heterogeneities along the elongated T cell body (dashed lines) as the nucleus completes one full back-and-forth oscillatory cycle.* See **Movie 10.**

**Supplemental Figure 2.**
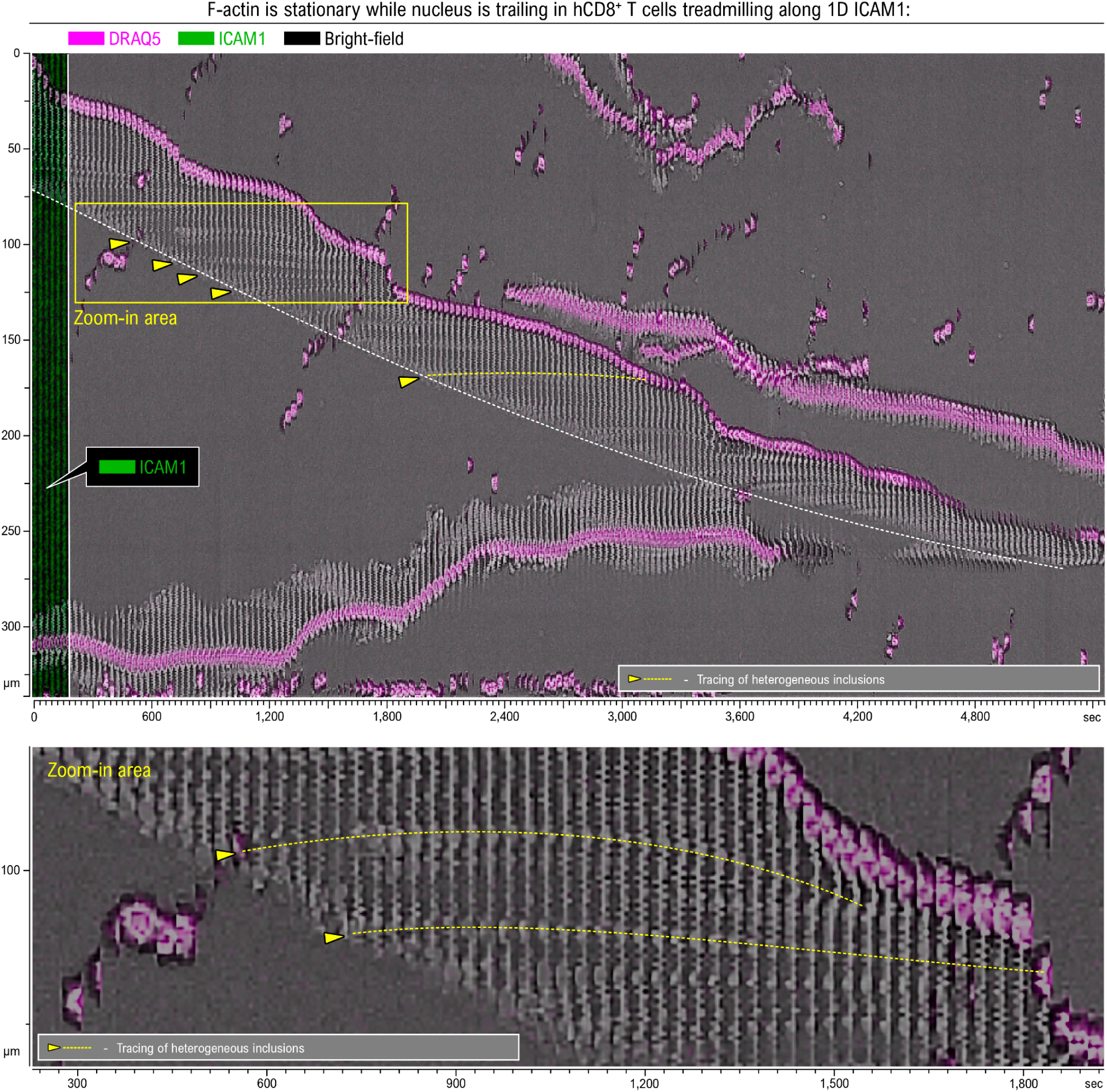
F-actin treadmilling within stationary protrusions as the nucleus trails during migration of a mature hCD8^+^ T cell on a 1 µm-wide ICAM1 lane. Kymograph of a migrating mature hCD8^+^ T cell showing stationary F-actin heterogeneities within elongated protrusions (dashed lines). The persistence of these stationary F-actin features, while protrusions extend along the ICAM1 lane and retract at the rear, is consistent with protrusion treadmilling as the nucleus trails behind the advancing leading edge. See **Movie 11.**

## MOVIE LEGENDS

**Movie 1. Population-scale amoeboid-to-mesenchymal mechanotype transition induced by septin inhibition in primary human hCD8**⁺ **T cells.** Human primary hCD8⁺ T cells migrating on orthogonal ICAM1 micropatterns initially display a compact amoeboid morphology under control conditions (+DMSO). Following addition of 50 µM UR214-9, numerous cells rapidly undergo a coordinated transition to an elongated mesenchymal mechanotype characterized by adhesion-guided spreading along ICAM1 lanes and the emergence of oscillatory nucleokinesis.

**Movie 2. Oscillatory nucleokinesis in multiple UR214-9-treated human primary hCD8**⁺ **T cells.** Human primary hCD8⁺ T cells with nuclei labeled by the live chromatin dye DRAQ5 (magenta) display oscillatory nuclear translocation after treatment with 50 µM UR214-9 while adhering to ICAM1 micropatterns. The movie illustrates that oscillatory nucleokinesis is a population-wide behavior rather than an isolated single-cell event.

**Movie 3. Oscillatory nuclear translocation in a UR214-9-treated primary human hCD8**⁺ **T cell.** The T cell is spread along a 1-µm-wide ICAM1 lane (green). The nucleus (DRAQ5, magenta) undergoes repeated back-and-forth translocations over ∼55 µm within an otherwise nearly stationary, elongated cell. Both brightfield and fluorescent channels are shown side-by-side.

**Movie 4. UR214-9-induced amoeboid-to-mesenchymal mechanotype transition in a primary human hCD8**⁺ **T cell.** Following addition of the septin GTPase inhibitor UR214-9, the T cell adheres and spreads along a 1-µm-wide ICAM1 lane (green), becomes highly elongated, and subsequently develops sustained oscillatory nucleokinesis.

**Movie 5. Stable oscillatory nucleokinesis in a UR214-9-treated primary human hCD8**⁺ **T cell.** Following the amoeboid-to-mesenchymal mechanotype transition, the elongated T cell remains stationary while the DRAQ5-labeled nucleus undergoes sustained, repeated back-and-forth translocations along the 1-µm-wide ICAM1 lane (green). Both brightfield and fluorescent channels are shown side-by-side.

**Movie 6. Acute dynein inhibition arrests oscillatory nucleokinesis in a UR214-9-treated primary human hCD8**⁺ **T cell.** The elongated T cell undergoes sustained oscillatory nucleokinesis until addition of the dynein inhibitor dynarrestin, which rapidly arrests nuclear translocation while the spread cell morphology remains largely unchanged until protrusions detach from the substrate and retract. Both brightfield and fluorescent channels are shown side-by-side.

**Movie 7. Dynein inhibition arrests oscillatory nucleokinesis independently of non-muscle myosin II activity.** A UR214-9-treated primary human hCD8⁺ T cell undergoes sequential treatment with para-nitroblebbistatin followed by dynarrestin during live-cell imaging. Oscillatory nucleokinesis persists after inhibition of non-muscle myosin II but becomes more irregular. Subsequent addition of dynarrestin rapidly arrests nuclear oscillation, after which the elongated cell slowly drifts in one direction. Brightfield, nucleus (DRAQ5), and both overlayed (with ICAM1) channels are shown side-by-side.

**Movie 8. Dynein inhibition arrests oscillatory nucleokinesis independently of non-muscle myosin II activity.** Another representative UR214-9-treated primary human hCD8⁺ T cell undergoes the same sequential treatment with para-nitroblebbistatin followed by dynarrestin during live-cell imaging. Oscillatory nucleokinesis persists after inhibition of non-muscle myosin II. Subsequent addition of dynarrestin rapidly arrests nuclear translocation, followed by collapse of protrusions and detachment of the cell. Brightfield, nucleus (DRAQ5), and both overlayed (with ICAM1) channels are shown side-by-side.

**Movie 9. Nuclear oscillation is decoupled from F-actin heterogeneity dynamics in an elongated UR214-9-treated human primary hCD8**⁺ **T cell on a 1-µm-wide ICAM1 lane.** During repeated oscillatory nuclear translocation, F-actin heterogeneities undergo slow centripetal displacement independently of nuclear motion, demonstrating the decoupling of nuclear oscillation from F-actin dynamics. Both brightfield and fluorescent channels are shown side-by-side.

**Movie 10. Nuclear oscillation is decoupled from F-actin heterogeneity dynamics in an elongated UR214-9-treated human primary hCD8**⁺ **T cell on a 2D ICAM1 grid.** During repeated oscillatory nuclear translocation, F-actin heterogeneities undergo slow centripetal displacement independently of nuclear motion, demonstrating the decoupling of nuclear oscillation from F-actin dynamics. Both brightfield and fluorescent channels are shown side-by-side.

**Movie 11. F-actin treadmilling within stationary protrusions of a migrating mature human primary hCD8**⁺ **T cell on a 1-µm-wide ICAM1 lane.** Stationary F-actin heterogeneities persist within elongated protrusions while the leading edge extends and the rear retracts, with the nucleus trailing behind the advancing protrusion, consistent with protrusion treadmilling.

**Movie 12. Alternating protrusion activation during oscillatory nucleokinesis in a UR214-9-treated primary human hCD8**⁺ **T cell.** The T cell is spread along a 1-µm-wide ICAM1 lane (green). The DRAQ5-labeled nucleus (magenta) undergoes repeated oscillatory translocations between opposing protrusions. Transient F-actin accumulation develops in the receiving protrusion before nuclear translocation and alternates between opposite protrusions prior to each reversal of nuclear movement. Both brightfield and fluorescent channels are shown side-by-side.

**Movie 13. Steady oscillatory nucleokinesis in a UR214-9-treated primary human hCD8**⁺ **T cell.** The T cell is spread along a 1-µm-wide ICAM1 lane (green). The DRAQ5-labeled nucleus (magenta) undergoes repeated back-and-forth translocations with an approximately constant velocity between turning points, corresponding to the steady oscillation pattern reproduced by the direct activation model. The left panel shows experimental data, while the right panel shows a simulation of the direct activation model.

**Movie 14. Accelerating oscillatory nucleokinesis in a UR214-9-treated primary human hCD8**⁺ **T cell.** The T cell is spread along a 1-µm-wide ICAM1 lane (green). The DRAQ5-labeled nucleus (magenta) undergoes repeated oscillatory translocations in which nuclear velocity progressively increases as the nucleus approaches the turning point, corresponding to the behavior reproduced by the delayed activation model. The left panel shows experimental data, while the right panel shows a simulation of the delayed activation model.

**Movie 15. Decelerating oscillatory nucleokinesis in a UR214-9-treated primary human hCD8**⁺ **T cell.** The T cell is spread along a 1-µm-wide ICAM1 lane (green). The DRAQ5-labeled nucleus (magenta) undergoes repeated oscillatory translocations in which nuclear velocity progressively decreases as the nucleus approaches the turning point, corresponding to the behavior reproduced by the direct activation with length-dependence model. The left panel shows experimental data, while the right panel shows a simulation of the direct activation with length-dependence model.

**Movie 16. Saltatory mesenchymal migration of UR214-9-treated primary human hCD8**⁺ **T cells.** The left panel shows an overlay of bright-field and fluorescence (DRAQ5-labeled nucleus: magenta; ICAM1 micropattern: green), while the right panel shows the DRAQ5 channel only. The cells undergo repeated oscillatory translocations between competing protrusions while the cell advances along the ICAM1 grid. Successive nuclear translocations resolve into net directional displacement through stride-wise cell migration.

**Movie 17. Dynein inhibition suppresses protrusion dynamics in tissue-resident zebrafish Langerhans cells.** Time-lapse imaging of zebrafish skin explants compares control conditions (+DMSO, top) with dynein inhibition by Dynapyrazole A (+DPA, bottom). Dynein inhibition markedly reduces protrusion extension-retraction dynamics and overall protrusion remodeling.

